# Asymmetric biparental but inefficient horizontal transmission of paralysis-causing sigmavirus in Queensland fruit fly

**DOI:** 10.64898/2026.03.07.710329

**Authors:** Sanjay Kumar Pradhan, Jennifer L. Morrow, Geraldine Tilden, Farzad Bidari, Shivanna Bynakal, Asokan Ramasamy, Markus Riegler

**Affiliations:** Hawkesbury Institute for the Environment, Western Sydney University, Locked Bag 1797, Penrith, NSW 2751, Australia; Department of Agricultural Entomology, University of Agricultural Sciences, Bengaluru - 560065, India; ICAR- Indian Institute of Horticultural Research, Hesaraghatta Lake (PO), Bengaluru - 560089, India

**Keywords:** Biparental transmission, CO_2_-triggered paralysis, insect-associated virus, negative-sense single-stranded RNA virus, rhabdovirus, RNA virus

## Abstract

Insects are associated with diverse RNA viruses, including vertically transmitted viruses that form persistent infections without apparent symptoms. One of the first documented vertically transmitted viruses is sigmavirus (*Rhabdoviridae*) affecting fitness of *Drosophila*. Sigmaviruses and related rhabdoviruses have also been detected in pest fruit flies and other arthropods. However, their prevalence, transmission, tissue localisation and fitness effects remain poorly known, despite their potentially common infections in diverse hosts. We investigated *Sigmavirus tryoni* (BtSV) prevalence, load, transmission across multiple generations and host effects in Queensland fruit fly (*Bactrocera tryoni*), Australia’s most significant horticultural pest, which carries BtSV at low prevalence (13.7%) across field populations. We detected BtSV in 6 of 12 laboratory populations (prevalence 12.5% to 80.4%) where it was transmitted biparentally within embryos. Although incomplete, maternal transmission was more reliable and resulted in higher BtSV load than paternal transmission. Paternally transmitted BtSV was almost entirely lost after two generations. BtSV became detectable in most uninfected individuals cohabiting with infected flies, but this resulted in a low load that was subsequently transmitted to only few offspring. BtSV occurred across developmental stages, digestive and reproductive tissues, albeit its viral load was lower in reproductive tissues when received paternally than maternally, and lower in testes than ovaries. Furthermore, BtSV-infected individuals suffered paralysis and mortality when exposed to high CO_2_ concentrations, a Rhabdoviridae effect previously reported for several *Drosophila* species, a muscid fly and mosquitoes. Our study suggests that sigmavirus transmission dynamics and fitness effects may apply broadly to arthropod hosts and affect their management.

## INTRODUCTION

Transcriptome analyses of insects have uncovered a vast diversity of RNA viruses (Shi et al. 2016). Many of these are predominantly found in insects and sometimes referred to as insect-specific (Carvalho and Long 2021; Li and Shi 2026) albeit host range of several insect-associated virus taxa also includes other arthropods, vertebrates and plants (Hogenhout et al. 2003). While the host effects of many RNA viruses detected in insect transcriptomes remain poorly characterised, some have substantial impacts. For example, several viruses contribute to colony collapse of honey bees (De Miranda and Genersch 2010), causing paralysis, thoracic and abdominal discolouration and hair loss, and death (Maori et al. 2007). Other viruses occur as infections that lack visible symptoms and are hence covert, yet can persist in insect populations and affect host fitness, physiology and behaviour (Fujiyuki et al. 2005; Han et al. 2015; Llopis-Giménez et al. 2017; Yuan et al. 2020; Hernández-Pelegrín et al. 2022), as well as influence vector competence of virus-transmitting insects (Talavera et al. 2018).

In general, viruses can be transmitted vertically, horizontally or a combination of both (Bézier et al. 2009). Vertical transmission typically occurs maternally from infected females to their offspring via the egg, or paternally via sperm (Morrow et al. 2023). Maternal transmission can occur within the egg (transovarial or transovarian) or on the surface of the egg (transovum) (Solter and Becnel 2017). In contrast, horizontal transmission can occur when individuals share resources that have been contaminated by infected individuals or when uninfected individuals interact with infected individuals by cohabitation or mating (Morrow et al. 2023).

One of the first described examples of a vertically transmitted virus was *Sigmavirus melanogaster* (Mononegavirales: *Rhabdoviridae*) (hereafter referred to as DmelSV), a single-stranded negative-sense RNA virus originally discovered in *Drosophila melanogaster* (L’Heritier and Teissier 1937; L’Heritier, 1970; Brun and Plus 1980; Longdon et al. 2011a). Diverse sigmaviruses and related rhabdoviruses have since been detected in other *Drosophila* species (Drosophilidae), true fruit flies (Tephritidae), house flies (Muscidae), sand flies (Psychodidae), black soldier flies (Stratiomyidae), a butterfly species (Nymphalidae), a fruit fly parasitoid wasp (Braconidae) and honey bees (Apidae), as well as in other arthropods such as *Varroa* mites (Laelapidae) (Fig. 1; Longdon et al. 2011b; Simmonds et al. 2016; Longdon et al. 2017; Remnant et al. 2017; Phumee et al. 2021; Sharpe et al. 2021; Pradhan et al. 2024a; Walt et al. 2024). Some rhabdoviruses such as the vesicular stomatitis virus (VSV) and the northern cereal mosaic virus also infect vertebrate animals and plants, respectively, and are transmitted by arthropods (Hogenhout et al. 2003). Detailed studies of transmission and host fitness effects have largely focussed on sigmaviruses associated with *Drosophila* (Fleuriet 1982; Longdon et al. 2011a), although vertical transmission has also been documented for sigmaviruses in other hosts including the Mediterranean fruit fly *Ceratitis capitata* and the butterfly *Pararge aegeria* (Longdon et al. 2015; Longdon et al. 2017), and for Diachasmimorpha longicaudata rhabdovirus (DlRhV) in the parasitoid wasp *Diachasmimorpha longicaudata* (Lawrence and Matos 2005). In *Drosophila,* DmelSV is reliably transmitted transovarially, with more virus received by the embryo maternally than paternally (Brun and Plus 1980; Longdon et al. 2012), albeit the relative importance of maternal versus paternal transmission has not been assessed for non-*Drosophila* hosts other than *C. capitata* and *P. aegeria* (Longdon et al. 2017). Rare horizontal transmission of DmelSV has also been detected, although the extent to which horizontally transmitted virus results in infection (Longdon and Jiggins 2012) and is transmitted to the next generation (L’Heritier 1948) is unclear. Tissue tropism studies in *Drosophila* show sigmavirus infection in all tissues including oesophagus, cardia, nerve ganglia, nerves and reproductive tissues (Bussereau 1970; Ammar et al. 2009), while DlRhV has been observed in eggs and the midgut lumen of first larval instars of *D. longicaudata*, as well as in the epidermis of parasitised *Anastrepha suspensa*, a tephritid fruit fly (Lawrence and Matos 2005). However, tissue tropism of sigmaviruses in other hosts remain uncharacterised.

**Fig. 1.**
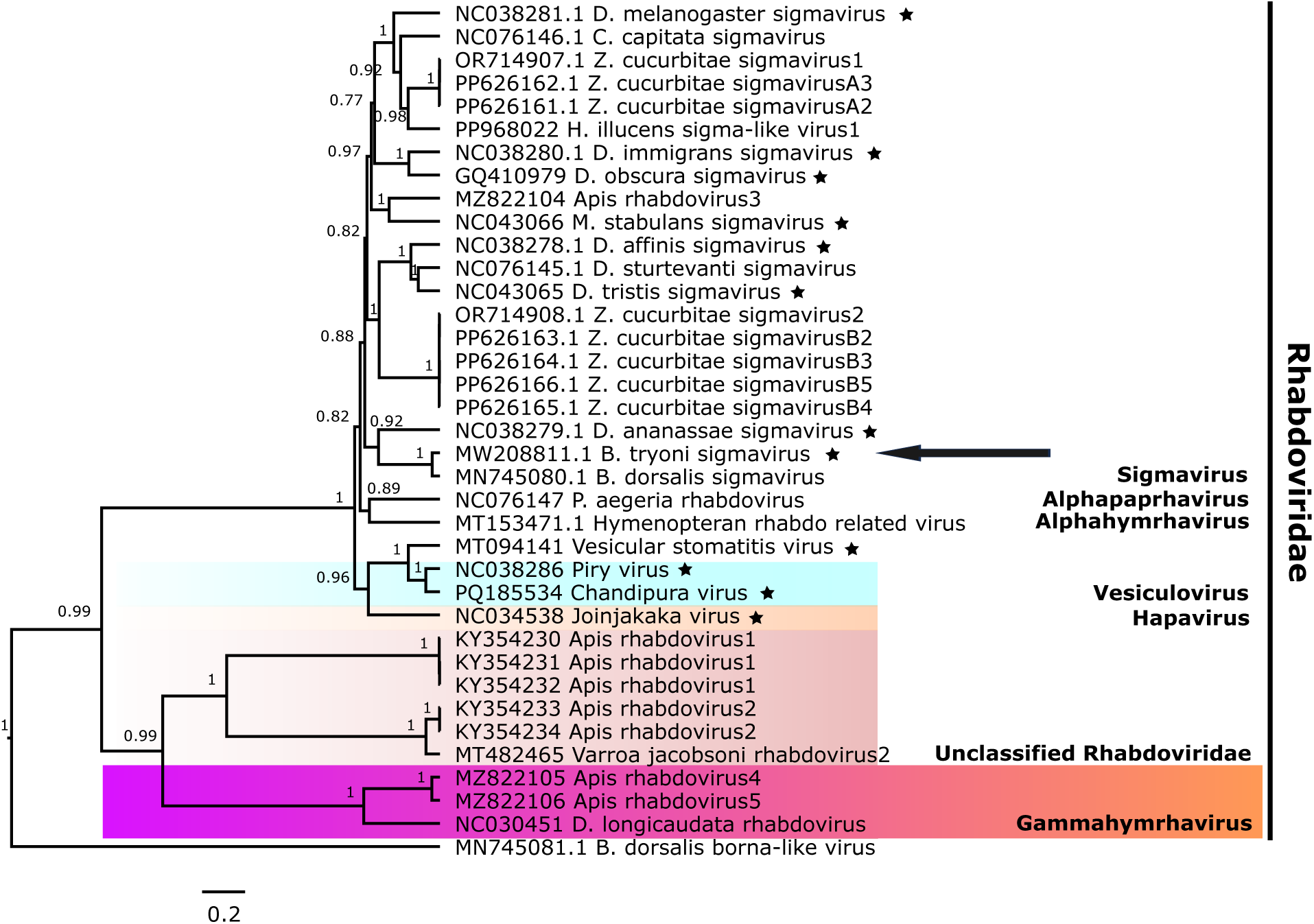
Phylogenetic tree of sigmaviruses and other *Rhabdoviridae*. The phylogenetic tree was constructed using Bayesian inference in Beast2 (parameters: birth–death model; 10,000,000 chain length; 25 % burn-in) of the RdRp (L) coding amino acid sequences of representatives of *Sigmavirus* (including *Sigmavirus tryoni* or Bactrocera tryoni sigmavirus BtSV, indicated by a horizontal arrow) and other *Rhabdoviridae* in Beast2. Conserved regions of the amino acid sequences were analysed using Gblocks. Posterior probability values are represented at the node, and values below 0.5 are not shown. The scale bar beneath the phylogenetic tree represents the number of amino acid substitutions per site. The stars indicate viruses that can cause CO_2_-triggered paralysis in hosts.

Without initially recognising a virus as the causative agent, L’Heritier and Teissier (1937) first observed that some *D. melanogaster* flies experienced irreversible paralysis and mortality after exposure to high concentrations of CO_2_ at cool temperatures whereas other flies fully recovered from the non-lethal stress of routine CO_2_ anaesthesia; this CO_2_-triggered paralysis was later linked to sigmavirus infection (L’Heritier, 1970; Brun and Plus 1980; Longdon et al. 2011a) causing nervous tissue damage after CO_2_ anaesthesia (L’Heritier 1948). As demonstrated for *D. melanogaster* experimentally infected with VSV, this lethality likely results from the virus glycoprotein (G) causing irreversible fusions of neurons and glia at low pH in the haemolymph due the exposure to high CO_2_ concentrations at cool temperatures; in contrast nitrogen (N_2_) anaesthesia did not trigger this paralysis (Chow et al. 2017).

The paralysis effect of sigmaviruses and related rhabdoviruses has been documented in multiple hosts beyond *Drosophila,* including the housefly *Muscina stabulans* and several mosquito species (Fig. 1; Rosen 1980; Longdon et al. 2012b). Besides this unusual CO_2_ effect, DmelSV infection was found to be costly to *D. melanogaster* by reducing the egg hatch rate (Seecof 1964; Fleuriet 1981a) and increasing developmental time and offspring mortality (Seecof 1964). Furthermore, at population level, DmelSV is thought to reduce survival of *D. melanogaster* in winter (Fleuriet 1981b). Conversely, under resource competition and crowded conditions, DmelSV replication in *D. melanogaster* is reduced (Yampolsky et al. 1999). However, relaxed selection can allow DmelSV persistence and spread despite its fitness costs. Furthermore, coevolutionary modelling shows alternating selective sweeps in host resistance and viral counteradaptation, driving shifts in infection prevalence with the virus typically leading the arms race (Wilfert and Jiggins 2013).

Sigmaviruses and rhabdoviruses with diverse genomes have also been found in five economically important tephritid fruit fly species. This includes the sigmaviruses found in the Queensland fruit fly *Bactrocera tryoni*, Australia’s most significant horticultural pest (Sharpe et al. 2021), *C. capitata* (Longdon et al. 2017), the oriental fruit fly *Bactrocera dorsalis* (Zhang et al. 2020) and the melon fly *Zeugodacus cucurbitae* (Pradhan et al. 2024a), as well as the rhabdovirus DlRhV detected in *A. suspensa* (Lawrence and Matos 2005). The genome structure of sigmaviruses of *Drosophila* species, *C. capitata* and *Z. cucurbitae*, as well as DlRhV of *D. laungicaudata* comprise six open reading frames (3’-N-P-X-M-G-L-5’), including the X ORF that encodes a protein of unknown function, whereas *Sigmavirus tryoni* of *B. tryoni* (hereafter BtSV) and the sigmaviruses of *B. dorsalis* and *P. aegeria*, as well as the rhabdoviruses detected in honey bees and *Varroa* mite do not have the X gene (Longdon et al. 2015; Simmonds et al. 2016; Remnant et al. 2017; Zhang et al. 2020; Sharpe et al. 2021; Pradhan et al. 2024a).

Previous research on the sigmaviruses of *C. capitata* and *P. aegeria* reported maternal and biparental transmission, respectively, based on multiple crosses and extensive offspring testing (Longdon et al. 2017). However, while sigmaviruses can cause persistent infections in laboratory populations (Longdon et al. 2017; Sharpe et al. 2021; Pradhan et al. 2024a), host effects of sigmaviruses in tephritids remain to be systematically evaluated. Persistent covert virus infections might negatively affect host fitness and performance, and thereby insect mass rearing (Maciel-Vergara and Ros 2017) and pest control strategies such as the sterile insect technique (SIT) used to control fruit fly and other pests (Llopis-Giménez et al. 2017; Sharpe et al. 2021). Conversely, insect-associated viruses could also constitute novel biocontrol agents (Coffman et al. 2024; Pradhan et al. 2024b; Coffman, 2025; Li and Shi 2025).

Besides BtSV, transcriptome analyses of *B. tryoni* laboratory populations detected Bactrocera tryoni cripavirus (*Dicistroviridae;* BtCV) and Bactrocera tryoni iflavirus (*Iflaviridae;* BtIV) (Sharpe et al. 2021), and Bactrocera tryoni orbivirus (*Sedoreoviridae*), Bactrocera tryoni toti-like virus (*Betatotivirineae*) and Bactrocera tryoni xinmovirus (*Xinmoviridae*) (Sharpe et al. 2025). A geographically broad survey of BtSV, BtCV and BtIV in field populations of *B. tryoni* from 29 sites across its distribution in Australia revealed that BtCV was the most prevalent virus (37.5% of 293 flies) followed by BtSV (13.7%) and BtIV (4.4%), whereby BtSV was detected in 22 of the 29 tested field populations (Sharpe et al. 2024). Furthermore, 46.8% of flies carried one virus while 4.8% were coinfected with BtCV and either BtSV or BtIV. However, BtSV did not coinfect any individuals with BtIV. It was hypothesised that this negative association pattern of BtSV and BtIV may be linked to their vertical transmission (Sharpe et al. 2024). In contrast to the horizontal transmission of BtCV, BtIV is maternally transmitted in *B. tryoni* (Morrow et al. 2023), and the sigmaviruses of *C. capitata* and other host species are vertically transmitted (Longdon et al. 2017), but this needs testing for BtSV in *B. tryoni*.

Here we comprehensively analysed the transmission pathways of BtSV in *B. tryoni*, evaluated its efficiency in maintaining persistent virus infections and tested whether BtSV induces CO_2_-triggered paralysis and mortality. This research builds on previous studies in other hosts and quantifies vertical (maternal and/or paternal) and horizontal transmission across multiple generations. We analysed the prevalence of BtSV in 12 *B. tryoni* laboratory populations, and then selected BtSV-infected and BtSV-free populations for transmission experiments. Then we analysed the BtSV load in different developmental stages, its localisation and the mode and efficiency of BtSV transmission across generations to evaluate the relative importance of maternal, paternal and horizontal transmission.

## MATERIALS AND METHODS

### Fly maintenance and virus screening

We used 12 *B. tryoni* laboratory populations (Supplementary Table S1). The populations were established with a single field-caught female (isofemale), few field-collected individuals or few laboratory-selected individuals >5 years prior to the experiments and then maintained as closed laboratory populations (6-8 generations per year). They can be considered inbred because previous research demonstrated that levels of heterozygosity and allelic richness of *B. tryoni* populations drop drastically after five generations in the laboratory (Gilchrist et al. 2012). The populations were kept at 25 °C, 60% relative humidity and natural light (Morrow et al. 2015). To prevent virus contamination, all cages and containers were washed with soap, then treated with household bleach (1%), and rinsed with water. Furthermore, fly cages (30 cm × 30 cm × 30 cm) were kept without any direct contact between cages (Morrow et al. 2023). From each population, total RNA was extracted from two pools each of five females and five males, in 500 µL TriSure (Meridian Biosciences) as per manufacturer’s instructions. cDNA was synthesised using the RevertAid First Strand cDNA synthesis kit (ThermoFisher) with a random hexamer and oligo d(T)-18 primer mix and screened by RT-PCR using virus-specific primers targeting the RNA-dependent RNA polymerase (RdRp) gene of *Sigmavirus tryoni* BtSV (variant BtRV1 MW208811) which has 34.6% amino acid similarity to DmelSV (Supplementary Table S2). cDNA quality was assessed using primers for *elongation factor 1 alpha* (*ef1a*), a host gene stably and highly expressed across all developmental stages of *B. tryoni* (Morrow et al. 2023). The RdRp amplicons of the populations lemon-eyes (LE+; + indicates a population is BtSV-infected whereby prevalence varied across populations) and C28+ were Sanger-sequenced to confirm virus variant identity. To assess BtSV prevalence, RNA was extracted from 12 females and 12 males of each laboratory population (28 females and 28 males of LE+), each in 500 µL TriSure, followed by cDNA synthesis and RT-PCR. We chose LE+ for the BtSV transmission, load and localisation experiments because it had the highest BtSV prevalence. B6– (– indicates BtSV-free population) and EC16– were used as the BtSV-free populations. All 12 populations were tested in the CO_2_ experiments.

### Characterisation of BtSV load across developmental stages

We provided mated LE+ females with a small cup containing larval diet covered with punctured parafilm for oviposition (Morrow et al. 2023) and collected eight embryos (24 hours old), eight third instar larvae (eight days after egg hatch), eight pupae (five days after pupation) and eight adults (four males and four females, five days after eclosion). RNA was extracted from each individual in 500 µL TriSure followed by cDNA synthesis. RT-qPCR was performed for two technical replicates using the BtSV-specific RdRp primers and *ef1a* primers, using SensiFast SYBR (Bioline) with the thermal cycling conditions of 95 °C for 3 minutes, then 45 cycles of 95 °C for 5 s, 60 °C for 10 s, and 72 °C for 20 s, followed by a disassociation cycle with an incremental increase of 1 °C from 50 °C to 98 °C every 5 s. RT-qPCR primers designed for BtSV RdRp and host *ef1a* (Supplementary Table S2) were tested for efficiency and detection limit using 10-fold dilution series; both primers exceeded 98% efficiency and the maximum Cq value for BtSV amplification under these conditions was 33 cycles. Thus, disassociation curves displaying a single temperature peak (±1.5 °C) and minimal primer-dimer noise combined with Cq values <33 were considered as virus-positive. RT-qPCR amplifications that did not meet these criteria were classified as virus-negative, with a quantification cycle (Cq) value assigned of 45, equal to the maximum number of cycles. All samples were normalised against transcripts of *ef1a*, using the formula 2^-(ΔCq)^ (Schmittgen and Livak 2008), which equated to a BtSV RdRp detection threshold of 10^-7^ per *ef1a* transcripts.

### BtSV transmission experiments

We conducted three types of transmission experiments: egg testing and surface bleaching experiments to assess whether BtSV transmission is vertical and, if so, whether it is transovarian (within the egg) or transovum (on the surface of the egg) (Supplementary Information 1a); crossing experiments to assess whether vertical BtSV transmission is maternal and/or paternal, and whether there is horizontal transmission between mating partners (Supplementary Information 1b); and single sex cohabitation experiments of infected and uninfected flies to assess horizontal transmission with a subsequent crossing experiment to assess vertical transmission of horizontally acquired BtSV ( Supplementary Information 1c). RNA was extracted from each individual in 500 µL TriSure, cDNA synthesised and subjected to RT-qPCR using *ef1a* as the normalisation gene. RT-qPCR melt curve analysis and Sanger sequencing of some RT-PCR amplicons were used to confirm BtSV identity.

For the embryo experiments (Supplementary Information 1a) we provided cages containing 100 LE+ females and males (21 days old) with an oviposition cup containing orange juice and covered with punctured parafilm for two hours. Embryos were collected and pools of 50 embryos were treated with household bleach (4% sodium hypochlorite) for 0, 1, 3 and 5 minutes in three replicates, followed by rinsing with deionised water.

For the crossing experiments to assess maternal, paternal and horizontal transmission between mating partners, (Supplementary Information 1b) we performed single-pair crosses each of LE+ × B6– (n= 10 pairs) and LE+ × EC16– (n= 8 pairs). For this, virgin females of unknown infection status were randomly collected from the LE+ population (with a BtSV prevalence of about 80%) and mated with virgin B6– or EC16– males in small containers provided with sugar, water and a sugar yeast hydrolysate mix (Morrow et al. 2015). After 14 days, the male was removed and stored at −80 °C while the female was provided with a small oviposition cup containing larval diet covered with punctured parafilm. As soon as the first offspring larvae left the larval diet and pupated, the adult females and their offspring pupae (five days old) were collected and stored at −80 °C. The BtSV load was then assessed for female, male and eight offspring pupae for each of the 18 pairs.

The remaining offspring of six high viral load LE+ females (families 1 to 6) were reared and separated into females and males upon emergence. For the maternal transmission assays this involved two offspring females each from families 1 and 2, and five offspring females each from families 5 and 6, individually crossed with a B6– male. For the paternal transmission assays, two offspring males each from families 1 to 4 and five offspring males each from families 5 and 6 were individually crossed with a virgin B6– female. From each pair that produced sufficient offspring, we collected the female, male and eight offspring pupae to assess their BtSV load. This allowed us to evaluate in a more controlled way than in the previous crossing experiment the maternal transmission (by 14 females) and paternal transmission (by 18 males) of BtSV, because all of these 32 individuals had received BtSV maternally (and not paternally or horizontally).

Then, the remaining offspring of 12 of these 32 individuals (two females each of matrilineal lines descending from females 1 and 2, and two males each of patrilineal lines descending from females 1 to 4) were reared and separated into females and males upon emergence and individually kept with B6– flies for 14 days. These crosses tested three transmission pathways: maternal-maternal-maternal (8 pairs), maternal-paternal-paternal (16 pairs), and maternal-paternal-maternal (15 pairs). Thus, we collected from 39 crosses the female, male and eight offspring pupae and assessed their BtSV load. This allowed us to evaluate BtSV transmission over three consecutive generations (starting with the high viral load LE+ females 1 to 6).

To assess transmission from high viral load grandmothers to a second offspring generation via maternal or paternal transmission, we caged 30 virgin female offspring of the two high viral load LE+ females 5 and 6 with 30 B6– males, and caged 30 virgin B6– females with 30 male offspring of the same two high viral load LE+ females. Embryos were collected from each of these four cages, and pools of 50 embryos each were bleached (4% sodium hypochlorite) for 0, 0.5, 1, 3 and 5 minutes before assessment of the BtSV load.

For the cohabitation experiments (Supplementary Information 1c), 15 virgin females (21 days old) each of LE+ and EC16– were placed in each of three cages, and 15 virgin males (21 days old) each of LE+ and EC16– in each of three other cages. All cages were supplied with water, sugar and sugar yeast hydrolysate mix (Morrow et al. 2015). LE+ adults have lemon-coloured eyes (Zhao et al. 2003) whereas EC16– flies have the wild-type red-brown eye colour. After one, five and ten days of cohabitation, five females and four males of BtSV-exposed EC16– of each cage were placed in microcentrifuge tubes and stored at −80 °C. The remaining ten females and eight males of BtSV-exposed EC16– of each cohabitation cage were individually kept in containers with water, sugar and sugar yeast hydrolysate mix for another five days (five females and four males) and ten days (five females and four males) of incubation after which the flies were also placed in microcentrifuge tubes and stored at −80 °C until all flies were tested for horizontally acquired BtSV.

Next we tested for any vertical transmission of horizontally acquired BtSV to the next generation. For this we kept 15 virgin female B6– flies with 15 female LE+ flies in one cage, and 15 virgin male B6– flies with 15 male LE+ flies in another cage for ten days, followed by ten days of incubation after removal of the LE+ flies. These BtSV-exposed virgin B6– flies were then individually mated with unexposed or exposed B6– flies and their offspring (8 pupae per family) were tested for BtSV along with the parents.

### BtSV localisation in adult LE+

BtSV load was quantified in dissected tissues (head, gut, ovaries and testes) of eight LE+ females and males, four offspring females and males each of high viral load LE+ females 5 and 6, four second generation matrilineal offspring females of high viral load LE+ females 1 and 2, and four second generation patrilineal offspring females and males of high viral load LE+ females 1 and 2. RNA was extracted from the head, gut and reproductive tissues in 200 µL TriSure for cDNA synthesis and BtSV load was quantified using virus-specific primers in RT-qPCR.

### CO_2_ exposure experiment

For the CO_2_ exposure experiment (Supplementary Information 1d) we collected 21 days old flies of the 12 laboratory populations, anaesthetised these with CO_2_ at room temperature for 30 s and placed them into small specimen jars (44 mm diameter, 55 mm height) covered with a mesh. For each population, five jars were filled with 20 flies (ten females and ten males). The flies quickly recovered from the short CO_2_ anaesthesia. The jars were then placed in a covered plastic container (7.5 L; 357 mm × 117 mm × 260 mm) which was placed into a temperature-controlled (12 °C) cabinet with an observation window. The container was filled with CO_2_ via a plastic tube for 2 minutes. Then the CO_2_ supply was stopped. The flies were kept in the CO_2_-filled container for 10 minutes followed by recovery at ambient air at 25 °C. Then, each cohort of 20 flies was maintained in a Bugdorm cage at room temperature with sugar, water and sugar yeast hydrolysate mix. As control, five replicates (each consisting of ten females and ten males) of each laboratory population were exposed to CO_2_ for 10 minutes at 25 °C, followed by recovery at ambient air at 25 °C. After CO_2_ exposure, mortality of the flies was recorded daily for 15 days. Dead flies were removed daily, and all surviving flies were collected after 15 days. RNA was extracted from all flies, cDNA synthesised and tested for BtSV by RT-PCR.

### Statistical analyses

The statistical analyses were performed using R (R Core Team 2020) with the packages multcomp (Hothorn et al. 2008) and ggplot2 (Wickham 2016). For viral load comparisons we used a non-parametric Kruskal–Wallis rank sum test that, if significant, was followed by a Dunn’s test adjusted for multiple comparisons using the Benjamini-Hochberg method. For the analysis of fly survival data, we fitted a Cox proportional hazards model using the survival package, and visualised survival curves in R with ggplot2 (Wickham 2016).

## RESULTS

### BtSV prevalence varied across laboratory populations

Six of the 12 laboratory populations (C28+, HAC+, GOS+, UNSW+, LE+ and OEWM+) were infected with BtSV (Supplementary Tables S1 and S3). BtSV had previously not been detected in C28+ (Sharpe et al. 2025), however Sanger sequencing of the RdRp PCR amplicons verified the BtSV presence in three of the 24 tested flies (Supplementary Information 2). In LE+, BtSV prevalence was 80.4% (Supplementary Table S1). RdRp amplicons from three individuals (one female, two males) were Sanger sequenced and matched BtSV (MW208811) over the region amplified by BtSV primer set 1 (Supplementary Information 2).

### BtSV load differed across developmental stages

BtSV load in LE+ varied significantly across developmental stages (Fig. 2a, Supplementary Information 3, Supplementary Table S4). The individual embryos (24 hours old) had substantially lower BtSV loads (∼3 logs) than pupae and adults (*p* = 0.001), however, they also yielded low total RNA with ∼10-cycle-higher Cq values for host *ef1a* than adult females (∼1000-fold lower template). Therefore, we interpreted this viral load result cautiously. Embryos and larvae were not sexed, so variation in viral load at these stages could partly reflect differences between males and females. Furthermore, larvae had an intermediate BtSV load, while pupae did not differ significantly from adults. Therefore, pupae were chosen to assess BtSV load of offspring in the maternal and paternal transmission experiments.

**Fig. 2.**
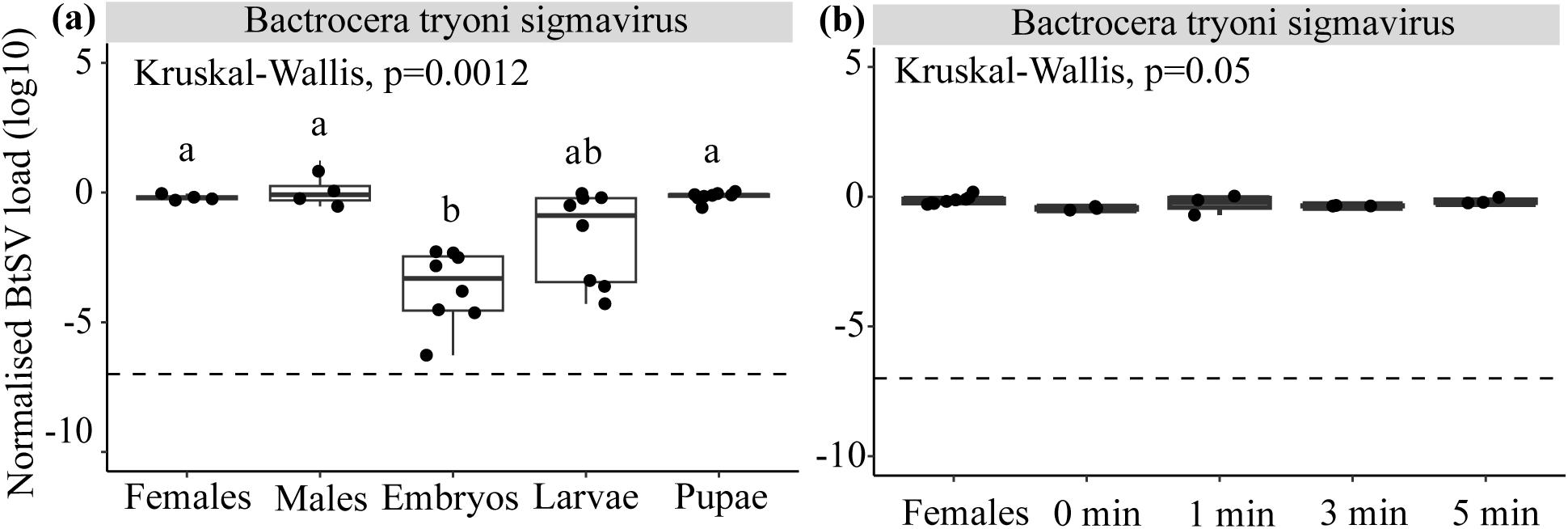
Bactrocera tryoni sigmavirus (BtSV) load across developmental stages and in embryos. **(a)** BtSV load in females, males, embryos, larvae and pupae of *Bactrocera tryoni* LE+ (four females, four males and eight individuals each for each other developmental stage). **(b)** Bactrocera tryoni sigmavirus (BtSV) load in *Bactrocera tryoni* LE+ females (n=8; mated with LE+ males) and embryos bleached for 0, 1, 3 and 5 minutes (three pools of 50 embryos each per treatment). Viral load (log_10_) was normalised to the transcript level of the host gene *ef1a*. The virus detection threshold was set at 10^-7^ BtSV per *ef1a* transcripts. Box plots show medians (centre line) and interquartile ranges (boxes). Significantly different pairwise comparisons identified by Dunn’s post hoc test are indicated by different letters.

### Embryo-bleaching reveals transovarian transmission of BtSV

The normalised BtSV load did not differ between LE+ females and pools of 50 embryos (<2 hours old) irrespective of embryo treatments with bleach (*p* = 0.05) (Fig. 2b, Supplementary Information 4, Supplementary Table S5). While single 24 hours old embryos had lower viral loads than adults (Fig. 2a) due to low RNA yield and ∼10-cycle-higher Cq values (∼1000-fold less template), pooling 50 embryos (<2 hours old; this section) yielded template levels comparable to females, possibly due to the differences in experimental procedures and/or *B. tryoni* generations used across the experiments. Overall, we expected that bleach treatment would remove surface-associated viral contamination, yet the persistence of similar viral loads before/after bleaching indicated that BtSV is transmitted vertically within the embryo rather than being on the external surface of the embryos.

### BtSV has both maternal and paternal transmission

Crosses between females with high BtSV load and uninfected males over three generations mostly produced offspring pupae with a similarly high viral load, indicative of maternal transmission. However, for some offspring, viral loads were more than 1000-fold lower than the maternal load, indicating imperfect maternal transmission. (Fig. 3a, b, and c; Supplementary Table S6, Supplementary Table S7).

**Fig. 3.**
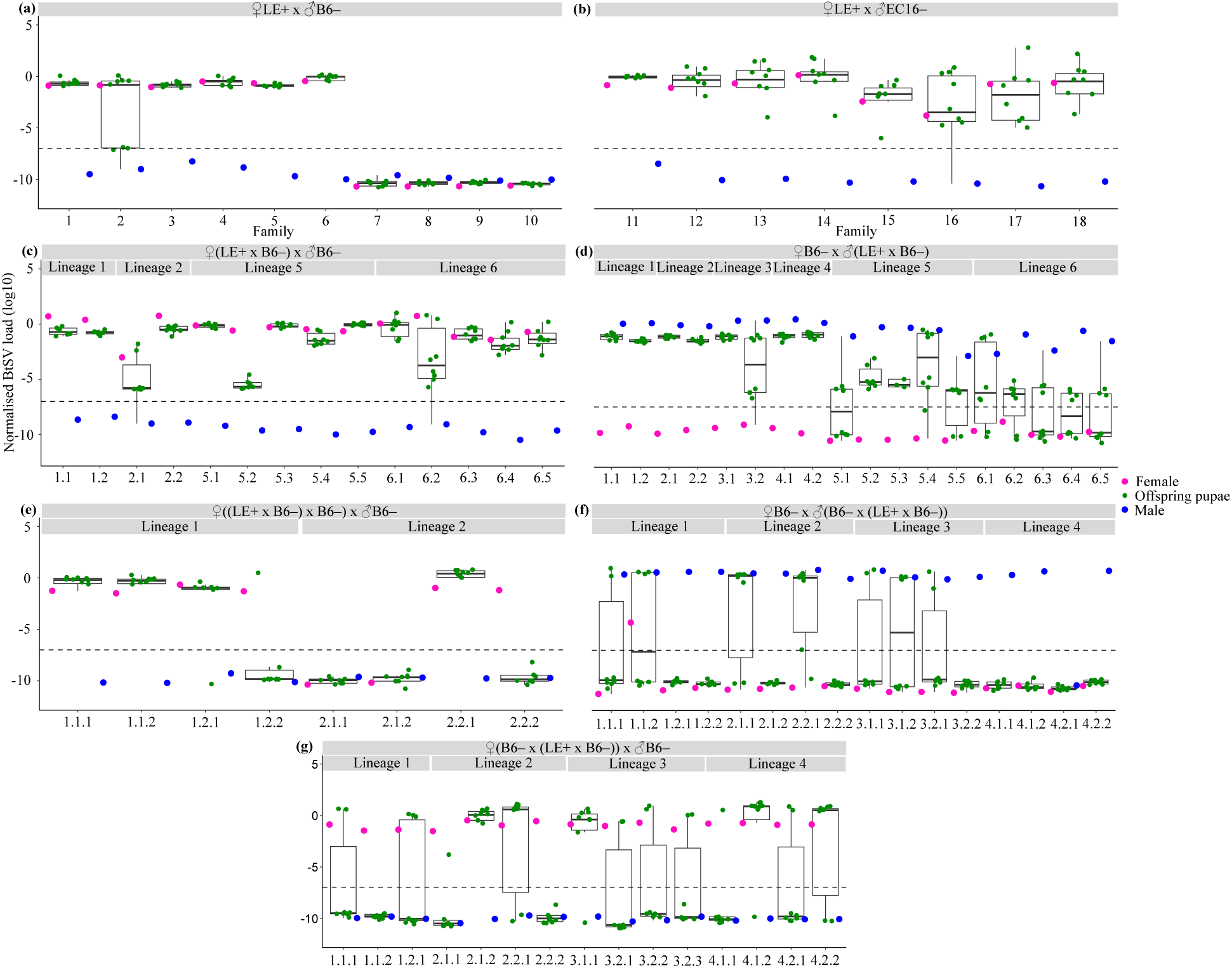
Vertical Bactrocera tryoni sigmavirus (BtSV) transmission across three generations. **(a)** BtSV load (log_10_) in ten LE+ females mated with BtSV-free B6– males and their offspring pupae (females pink dots on left; males blue dot on right; eight offspring pupae green dots in centre); **(b)** eight LE+ females mated with BtSV-free EC16– males and their offspring pupae; **(c)** two offspring females each of females 1 and 2, and five offspring females each of females 5 and 6, all mated with BtSV-free B6– males, and eight of their offspring pupae; **(d)** two offspring males each of females 1 to 4, and five offspring males each of females 5 and 6, all mated with BtSV-free B6– females, and eight of their offspring pupae; **(e)** two females each of matrilineal lines descending from females 1 and 2, all mated with BtSV-free B6– males, and eight of their offspring pupae; **(f)** two males each of patrilineal lines descending from females 1 to 4, all mated with BtSV-free B6– females, and eight of their offspring pupae; **(g)** two females each of matri-patrilineal lines descending from females 1 to 4, all mated with BtSV-free B6– males, and eight of their offspring pupae. BtSV load was normalised to the transcript level of the host reference gene *ef1a*. BtSV detection threshold was set at 10^-7^ per *ef1a* transcripts (dashed line). Points below this threshold represent tested samples without valid detectable amplification. Box plots show medians (centre line) and interquartile ranges (boxes). No vertical jitter was applied to the y-axis.

In comparison, 18 males that had inherited a high BtSV load from infected mothers, mated with BtSV-free virgin B6– females also had BtSV infected offspring pupae. While offspring pupae for eight of these males had a consistently high BtSV load, the BtSV load in the offspring pupae of the remaining ten males was lower or absent (Fig. 3d; Supplementary Table S7), suggesting that paternal transmission is less reliable than the maternal transmission.

### Maternal transmission, although imperfect, is more efficient than paternal transmission

We then assessed BtSV transmission in 39 individuals that had received BtSV either through maternal transmission only (matrilineal) or a mix of maternal and paternal transmission (matri-patrilineal) over consecutive generations. For the maternal-maternal-maternal transmission pathway (eight females), four females with a high BtSV load produced offspring pupae with a similarly high BtSV load, two females with a high BtSV load produced offspring pupae mostly without detectable BtSV load, and two females without BtSV produced offspring pupae without BtSV (Fig. 3e, Supplementary Table S7).

For the maternal-paternal-paternal transmission pathway (16 males), all males had a high BtSV load, yet their offspring pupae had no detectable BtSV, except for a few pupae of seven males (Fig. 3f, Supplementary Table S7). This loss of BtSV was less drastic for the maternal-paternal-maternal transmission pathway (15 females, all with high BtSV load) where BtSV was still detected in some offspring pupae of 13 females (Fig. 3g, Supplementary Table S7).

Embryo analyses confirmed these trends: females that had received BtSV from females 5 and 6 and mated with BtSV-free B6– males, produced embryos with a BtSV load as high as in these females, irrespective of embryo treatments with bleach (*p* =0.75, *p* =0.46) (Fig. 4a, Supplementary Information 5, Supplementary Table S8). In contrast, males that had received BtSV from females 5 and 6, and mated with BtSV-free B6– females, fathered embryos with a significantly lower BtSV load, irrespective of embryo treatments with bleach (*p* = 0.01; *p* = 0.009) (Fig. 4b, Supplementary Information 5, Supplementary Table S8).

**Fig. 4.**
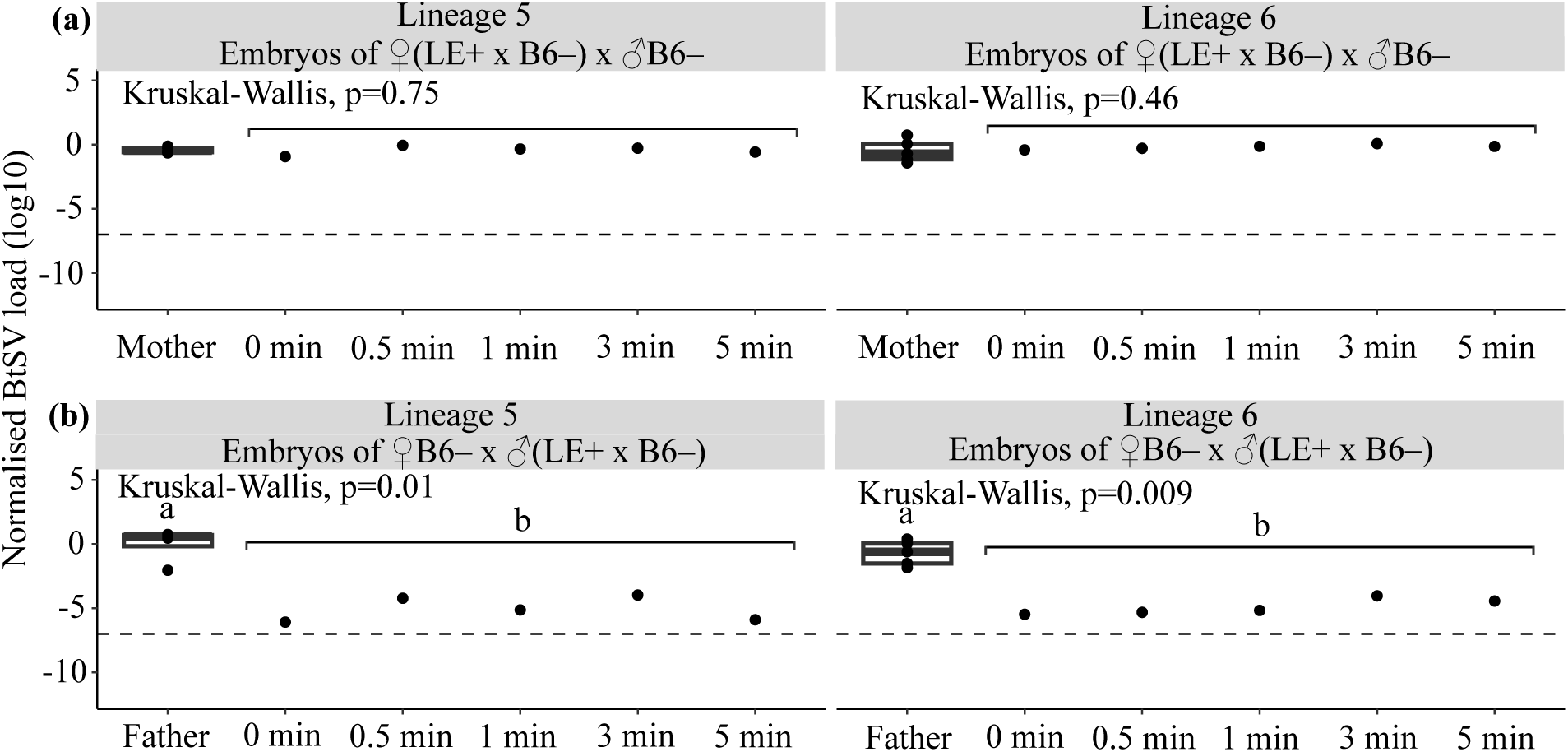
Bactrocera tryoni sigmavirus (BtSV) load in embryos. BtSV load (log_10_) in pooled embryos (50 embryos each, bleached for 0, 0.5, 1, 3 and 5 minutes) of **(a)** 30 female offspring of LE+ females 5 and 6 kept in one cage with BtSV-free 30 B6– males and **(b)** 30 male offspring of LE+ females 5 and 6 kept in a cage with 30 BtSV-free B6– females. BtSV load was normalised to the transcript level of the host reference gene *ef1a*. BtSV detection threshold was set at 10^-7^ *ef1a* transcripts. Box plots show medians (centre line) and interquartile ranges (boxes). Significantly different pairwise comparisons identified by Dunn’s post hoc test are indicated by different letters.

### Flies can horizontally acquire BtSV but it is not efficiently transmitted to the next generation

After cohabitation with LE+ females, we detected BtSV in most exposed females of the uninfected EC16– population, except for four females with one day cohabitation followed by ten days of incubation (1:10; Fig. 5a). Similarly, after cohabitation with LE+ males, we detected BtSV in most exposed males of the uninfected EC16– population, except for three males with one day cohabitation followed by ten days of incubation (1:10; Fig. 5b). For both sexes, horizontally acquired BtSV loads in EC16– flies were markedly lower than those observed in LE+ flies, but due to variability among individuals and limited sample sizes, these differences only reached statistical significance for two female combinations (1:10 and 5:5; *p* = 0.0005) (Fig. 5a, Supplementary Information 6, Supplementary Table S9), and for one male combination (1:10; *p* = 0.0008) (Fig. 5b, Supplementary Information 6, Supplementary Table S9). Furthermore, there was no significant difference in the BtSV load between EC16– females and males (*p* = 0.28) that had horizontally acquired BtSV from cohabiting LE+ individuals (Supplementary Information 6).

**Fig. 5.**
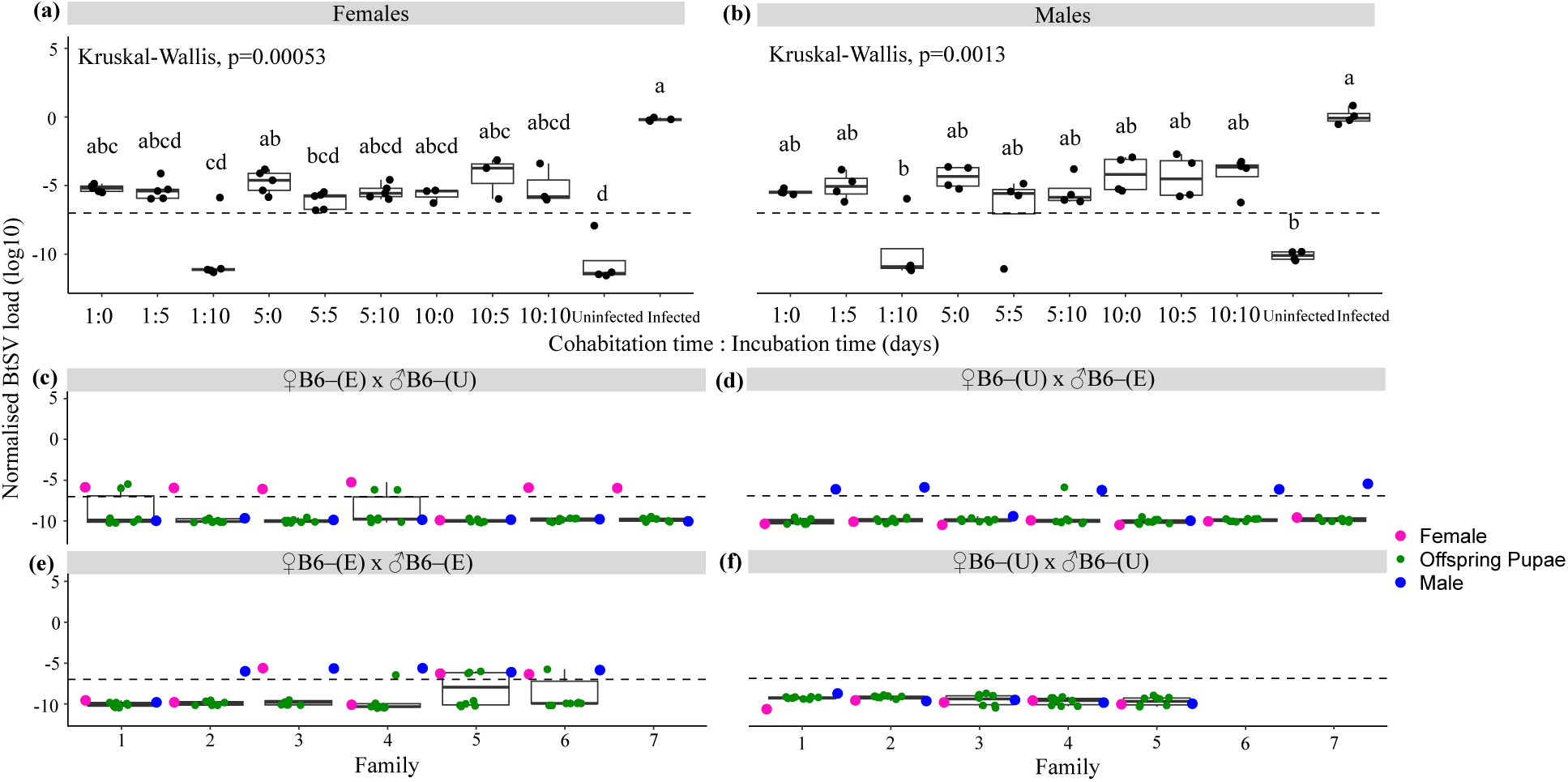
Horizontal Bactrocera tryoni sigmavirus (BtSV) transmission. BtSV load (log_10_) in originally BtSV-free EC16– **(a)** females and **(b)** males cohabiting with LE+ females and males, respectively for 1, 5 and 10 days followed by additional incubation in isolation for 0, 5 and 10 days. BtSV load (log_10_) in BtSV-free B6– females (pink dots on left) and males (blue dots on right) first exposed (E) to LE+ flies by cohabitation (10 days), followed by additional incubation (10 days) and then mated with unexposed (U) BtSV-free B6– flies in four combinations: **(c)** B6–(E) × B6–(U); **(d)** B6–(U) × B6–(E); **(e)** B6–(E) × B6–(E); **(f)** B6–(U) × B6–(U) (control). The BtSV load of their offspring pupae (n = 8) is presented as green dots in centre. BtSV load was normalised to the transcript level of the host reference gene *ef1a*. BtSV detection threshold was set at 10^-7^ *ef1a* transcripts. Box plots show medians (centre line) and interquartile ranges (boxes). Significantly different pairwise comparisons identified by Dunn’s post hoc test are indicated by different letters.

However, horizontally acquired BtSV was rarely transmitted to the next generation. Of 56 offspring pupae of BtSV-exposed B6– females mated with B6– males, only two pupae each from two of the seven BtSV-exposed females had BtSV, all at a low virus load (Fig. 5c). In the reciprocal cross, of 56 tested pupae, only one offspring pupa from one of the seven BtSV-exposed males had BtSV, also at a low virus load (Fig. 5d). Of the 48 tested pupae produced by the six pairs of BtSV-exposed females and males, a low BtSV load was detected in five offspring pupae of three BtSV-exposed pairs; for two of these pairs both the female and the male had acquired BtSV (Fig. 5e, Supplementary Table S10). In total this vertical transmission of horizontally acquired BtSV occurred in 10 of 160 tested pupae (6.25%). In comparison, no BtSV was detected in the control crosses and their offspring (Fig. 5f).

### BtSV did not show tissue tropism in adult LE+

For individuals randomly taken from the LE+ population, BtSV load did not differ significantly in female tissues (head, gut and ovaries) (*p* = 0.5) and male tissues (head, gut and testes) (*p* = 0.88) (Fig. 6a, Supplementary Information 7, Supplementary Table S11). However, the BtSV load was highly variable, probably because we tested individuals for which we did not know whether they had received BtSV maternally, paternally or horizontally.

**Fig. 6.**
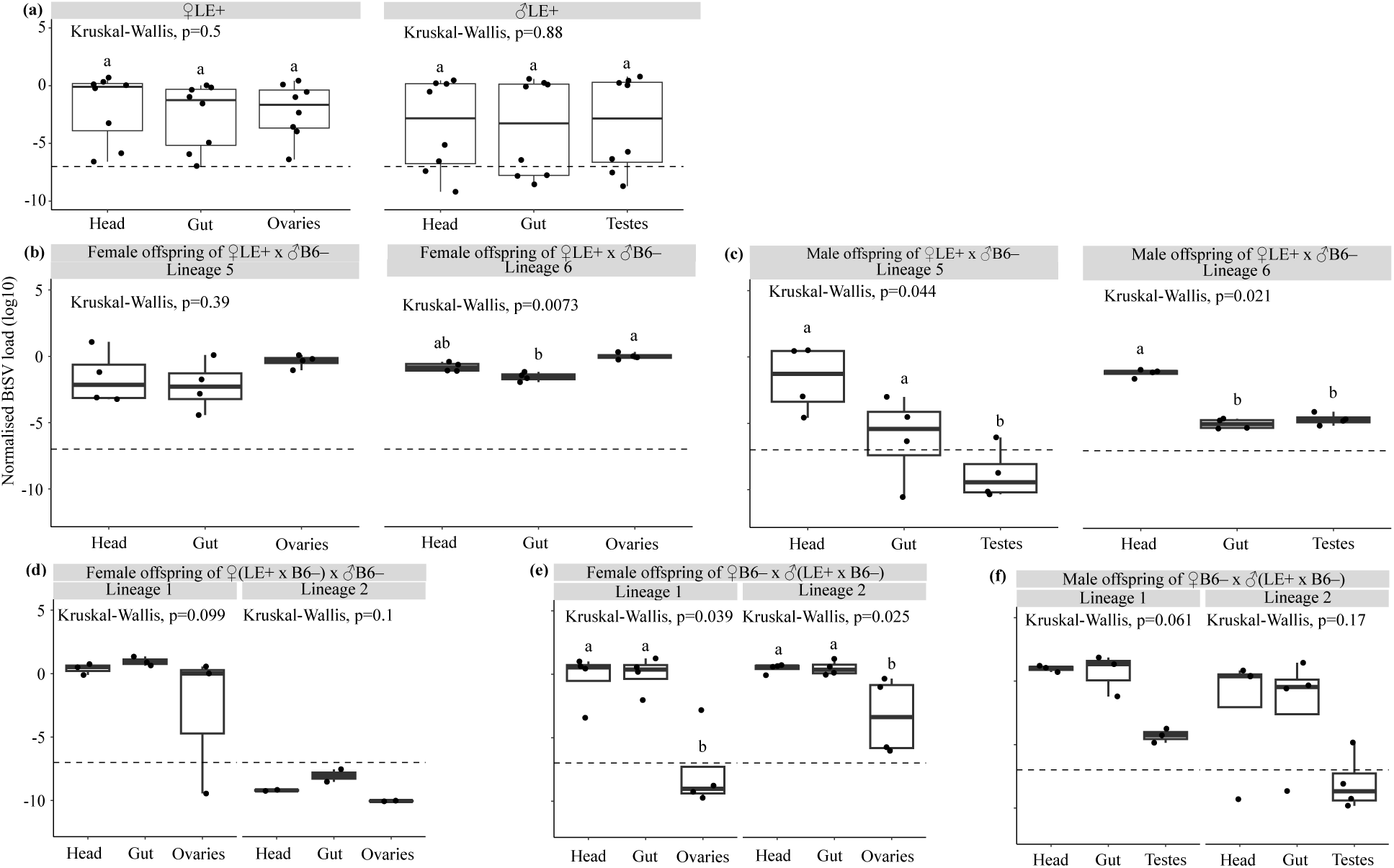
Bactrocera tryoni sigmavirus (BtSV) tissue tropism in females, males and their offspring. **(a)** BtSV load (log_10_) in tissues of 21 day old LE+ females (head, gut and ovaries) and males (head, gut and testes) (n=8 for each sex); **(b)** female offspring (head, gut and ovaries; n = 4) of high viral load females 5 and 6; **(c)** male offspring (head, gut and testes; n = 4) of high viral load females 5 and 6; **(d)** female offspring (head, gut and ovaries; n = 4) of matrilineal lineages descending of high viral load females 1 and 2; **(e)** female offspring (head, gut and ovaries; n = 4) of patrilineal lineages descending of high viral load females 1 and 2; **(f)** male offspring (head, gut, and testes; n = 4) of patrilineal lineages descending of high viral load females 1 and 2. BtSV load was normalised to the transcript level of the host reference gene *ef1a*. BtSV detection threshold was set at 10^-7^ *ef1a* transcripts. Box plots show medians (centre line) and interquartile ranges (boxes). Significantly different pairwise comparisons identified by Dunn’s post hoc test are indicated by different letters.

Therefore, we tested the BtSV load in head, gut and ovaries of female offspring of the two high viral load LE+ females 5 and 6. Overall, the virus load across the tissues was less variable than in the previous experiment. Furthermore, we found that it did not significantly differ in tissues for offspring of female 5 (*p* = 0.39), whereas for offspring of female 6, ovaries had significantly higher BtSV load than the gut (*p* = 0.007) with the head not being different from these two tissues (Fig. 6b, Supplementary Information 7, Supplementary Table S11). In contrast, in male offspring of female 5, head and gut had significantly higher BtSV load than testes (*p* = 0.04), whereas in male offspring of female 6, the head had significantly higher BtSV load than gut and testes (*p* = 0.02) (Fig. 6c, Supplementary Information 7, Supplementary Table S11).

We further analysed BtSV load in tissues of female second-generation offspring of high BtSV load LE+ females 1 and 2 (maternal-maternal transmission pathway) and found no significant difference in the female tissues (*p* = 0.099, *p* = 0.1) albeit virus had been lost entirely in one lineage (Fig. 6d, Supplementary Table S11). For the tissues of male and female second-generation offspring of high BtSV load females 1 and 2 (maternal-paternal transmission pathway) we found that females had lower BtSV load in ovaries than head and gut (*p* = 0.04, *p* = 0.02) (Fig. 6e) whereas males also had lower BtSV load in testes than head and gut (*p* = 0.06, *p* = 0.17) (Fig. 6f, Supplementary Table S11).

### CO_2_-induced paralysis and mortality in BtSV-infected fly populations

Flies of all BtSV-infected populations experienced moderate to severe paralysis after CO_2_ was applied at 12 °C for ten minutes. Paralysed flies displayed uncoordinated movement, and while some flies started moving and flying again, they appeared less active than flies of the uninfected populations. Within 15 days, BtSV-infected populations experienced very high levels (up to 90%) mortality (Fig. 7a) and most of the dead flies had a high BtSV load (Supplementary Information 8, Supplementary Table S12). Exceptions to this were the OEWM+ and C28+ populations which had lower BtSV load (Supplementary Information 8) and lower levels of mortality (∼25% and ∼15% after 15 days, respectively), but both also had a lower BtSV prevalence of ∼21% and 12.5%, respectively (Supplementary Table S1). Mortality of flies of BtSV-infected populations mostly occurred within 24 to 72 hours after CO_2_ treatment at 12 °C. In contrast, flies of BtSV-free populations fully recovered after CO_2_ treatment, and did not experience high levels of mortality (<10%) within 15 days after CO_2_ exposure at 12 °C. The surviving flies were mostly BtSV-free, albeit a few surviving flies showed weak PCR amplicons (Supplementary Information 9). Survival analysis revealed that BtSV-infected populations had a significantly higher mortality risk than uninfected populations (Cox proportional hazards model: hazard ratio = 1.89, 95% CI= 1.33-2.69, *p*= 0.00039; Supplementary Information 10). In the control experiment (exposure to CO_2_ at 25 °C for ten minutes), mortality in all fly populations was also low (<10%) (Fig. 7b, Supplementary Table S13).

**Fig. 7.**
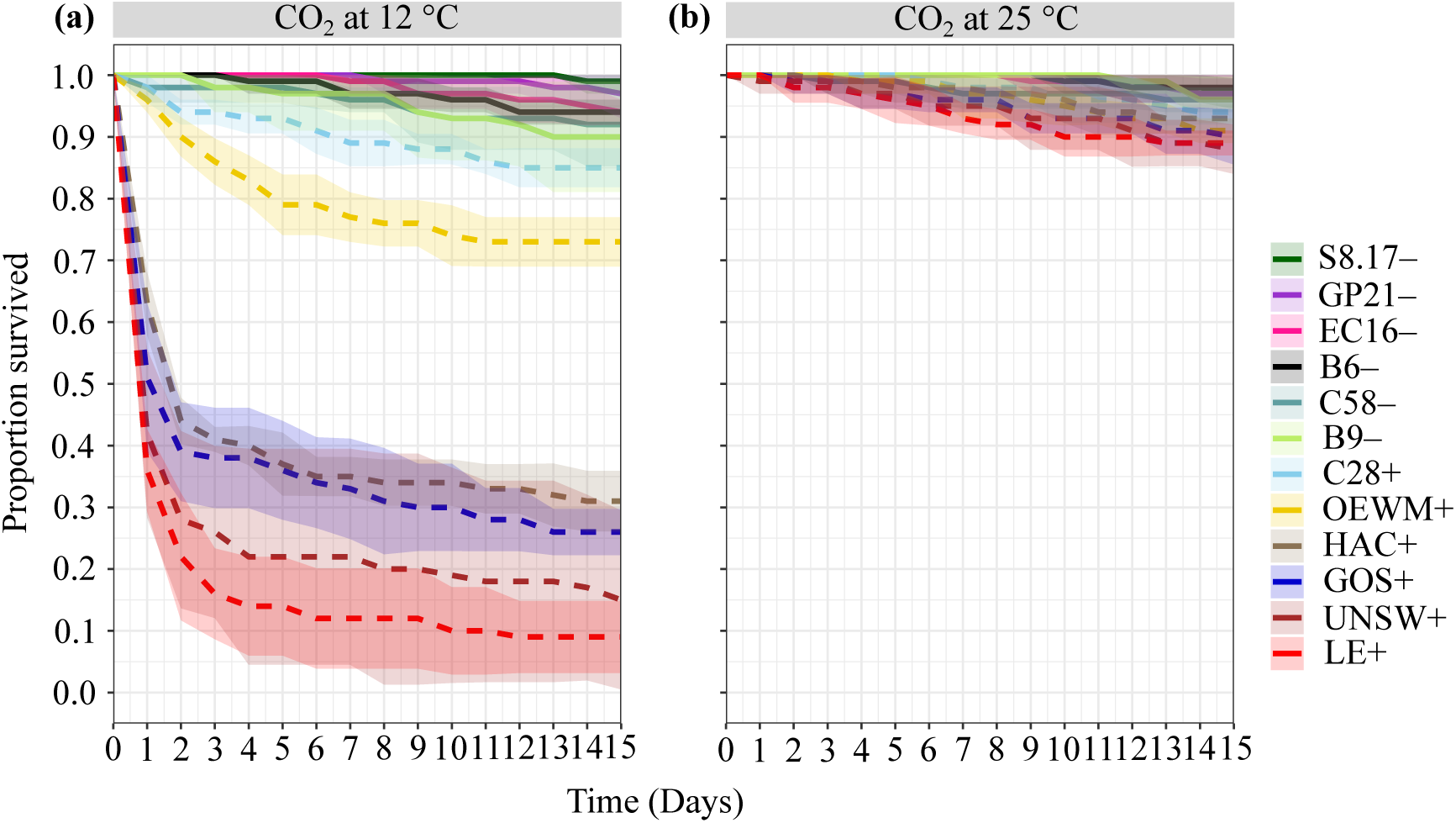
CO_2_-triggered paralysis in Bactrocera tryoni sigmavirus (BtSV)-infected *Bactrocera tryoni*. **(a)** Fly survival in 12 *Bactrocera tryoni* laboratory populations exposed to CO_2_ at 12 °C for 10 minutes. **(b)** Fly survival in the same populations exposed to CO_2_ at 25 °C for 10 minutes. BtSV-infected populations (+) are represented by dashed lines, and BtSV free populations (–) are represented by continuous lines. The standard error is represented by the shaded ribbon around the lines.

## DISCUSSION

Our study characterised BtSV prevalence, transmission pathways, developmental stage and tissue tropism, as well as CO_2_-triggered paralysis and mortality in laboratory populations of *B. tryoni*, an insect pest of economic significance. We detected BtSV in 6 of 12 laboratory *B. tryoni* populations at prevalences between 12.5% and 80.4%, including populations in which BtSV was more prevalent than a previous study of field populations (Sharpe et al. 2024). This previous study reported BtSV in 40 of 293 individuals (13.65%) collected from 29 field populations across Australia, suggesting that BtSV may either be costly, or environmental and/or genetic factors may limit BtSV infections. BtSV infected all developmental stages of *B. tryoni*, with the highest viral load occurring in pupae and adults. BtSV was vertically transmitted within embryos, via both females and males, whereby maternal transmission was more reliable than paternal transmission. We also detected horizontal transmission but this resulted in a low BtSV load that was not efficiently transmitted to the next generation. BtSV was found to infect nervous, digestive and reproductive tissues in both sexes equally. Furthermore, BtSV-infected flies suffered paralysis and high mortality when exposed to CO_2_ at 12 °C for 10 minutes whereas uninfected individuals recovered after treatment. The paralysis effect was not seen when flies were exposed to CO_2_ at 25 °C. This CO_2_-triggered sigmavirus effect has so far been revealed for *Drosophila* species, the housefly *M. stabulans* (Longdon et al. 2011b), and CO_2_-triggered paralysis was also recorded for other rhabdoviruses in *Drosophila* and mosquitoes (Bussereau 1975; Rosen 1980). Therefore, this effect may apply to infections of arthropods with sigmavirus and related rhabdoviruses more widely (Chow et al. 2017).

### Biparental BtSV transmission

RNA viruses in insects spread by vertical or horizontal transmission, or a mixture of both. Using a detailed assessment and various experimental approaches we found that BtSV has within-embryo transmission in *B. tryoni*, with transmission from both females and males to their offspring. This is similar to DMelSV transmission in *D. melanogaster* (Longdon and Jiggins 2012) where high transmission efficiency relies on early infection of germline cells during development (Fleuriet 1988). DMelSV is transmitted via both eggs and sperm (Longdon et al. 2010) and persists in populations despite fitness costs (Fleuriet 1981a). However, females transmit the virus more efficiently compared to males. Yet, if a female acquires DMelSV paternally, the transmission rate to the next generation drops from 98% to 20%. In contrast, males fail to transmit DMelSV to consecutive generations if they receive it paternally with transmission dropping from 45% to 0% (Longdon et al. 2011a). We found for BtSV in *B. tryoni* that maternal transmission remained efficient across generations, resulting in high viral load, whereas paternal transmission was less efficient across consecutive generations. Notably, paternal transmission to the second generation was still possible, even resulting in a high BtSV load in some individuals but at an overall low transmission efficiency (∼20%), whereas female transmission resulted in a high BtSV load in some individuals at a moderate transmission efficiency (∼40%). The reduced efficiency of paternal transmission may reflect a dilution of BtSV load because sperm may deliver less virus to embryos compared to eggs. This is supported by findings of the embryo bleaching experiments (showing lower viral loads in paternally infected embryos) and the tissue localisation analyses (revealing lower viral loads in testes than ovaries).

### Inefficient horizontal BtSV transmission

Horizontal transmission of viruses such as BtCV in *B. tryoni* (Morrow et al. 2023) and nora virus in *D. melanogaster* (Ekström and Hultmark 2016) likely occurs through food sharing, and interactions of flies with infected individuals through regurgitation and defecation. Our results show that BtSV-infected flies can transmit BtSV horizontally via cohabitation but this results in low viral loads. Such horizontal acquisition of virus may not result in infection, be transient and restricted to the gut resulting in no or limited transmission to the next generation. We found that only 10 out of 160 offspring (6.25%) of individuals that had become BtSV-positive after cohabitation with infected flies were BtSV positive suggesting that the horizontal-vertical transmission pathway is inefficient. Furthermore, BtSV was more frequently detected in uninfected flies exposed to groups of infected flies as part of the single sex cohabitation experiments (15 infected and 15 uninfected individuals cohabiting in a cage for up to 10 days) whereas BtSV did not become detectable in uninfected flies of the single pair crossing experiments (individual pairs of infected and uninfected flies kept together in a container for 14 days). Therefore, fly density may play an important role in horizontal transmission, whereby fly densities are likely higher in laboratory settings than the field (Sharpe et al. 2021).

Host genotype can affect susceptibility, replication and transmission efficiency of insect-associated viruses (Imrie et al. 2021). For the BtSV transmission experiment we used one infected (LE+; 80% BtSV prevalence) and two uninfected (B6– and EC16–) populations. Albeit all three can be considered highly inbreed, they may differ in genetic factors relevant to BtSV transmission.

### CO_2_-triggered paralysis and mortality

BtSV caused paralysis and mortality in *B. tryoni* exposed to high CO_2_ concentrations at 12 °C. Such a CO_2_-triggered paralysis caused by sigmaviruses was also reported for *Drosophila* and *M. stabulans* (Longdon et al. 2011a, b). However, the BtSV effect in *B. tryoni* was different from that observed for DmelSV in *Drosophila*: the paralysis was temporary, and flies showed signs of recovery 2-3 hours after the treatment; yet very high levels of mortality were observed in these flies after 24 hours. In contrast, DmelSV caused irreversible paralysis in *Drosophila* that led to mortality within 30 minutes (Longdon et al. 2011a, b). In a few surviving *B. tryoni* flies, weak BtSV PCR amplicons were observed, as also seen by Wilfert and Jiggins (2010). This suggests a threshold viral load may be required to cause a CO_2_ effect, or detection of non-infectious sigmavirus. This needs further investigation, together with assessment of the effects of horizontally acquired BtSV in *B. tryoni*. In *Drosophila* no CO_2_-triggered paralysis was observed in mating partners of infected flies and it was concluded that horizontal transmission is very rare (Longdon et al. 2011a).

Besides sigmavirus, other Rhabdoviridae have been shown to cause CO_2_-triggered paralysis in *Drosophila*: including VSV (Chow et al. 2017), piry virus and chandipura virus (Bussereau 1975); all three infecting various hosts including mammals, ticks and insects. CO_2_-triggered paralysis by some rhabdoviruses has also been reported for the mosquitoes *Toxorhynchites amboinensis, Culex quinquefasciatus* and *Aedes albopictus* (Rosen 1980). Therefore, it is likely that sigmaviruses and other rhabdoviruses such as those of honey bee and *Varroa* (Remnant et al. 2017; Li et al. 2023) may also cause CO_2_-triggered paralysis, with potential implications for bee keeping and *Varroa* management.

Previous studies tested whether exposing tephritid-infested fruit to high CO_2_ concentrations, including at cool temperatures, would lead to fruit fly mortality and hence disinfestation of infested fruit. Alonso et al. (2005) found that treating mandarin fruit infested with *C. capitata* larvae with 95% CO_2_ at 20 and 25 °C for 20 hours resulted in high larval mortality without affecting fruit quality. Similarly, Golding et al. (2010), showed that *B. tryoni* larvae in peaches and cherries were killed when exposed to 95% CO_2_ at 0 °C for 24 to 48 hours before ambient storage, effectively disinfesting fruit without quality loss. Exposure to high CO_2_ concentrations for 1 or 2 days as in these two studies might be effective in killing the flies in the absence of sigmavirus. However, the sigmavirus infection status of *C. capitata* and *B. tryoni* used in these disinfestation experiments is unknown, yet its presence could potentially have increased fly mortality. It is also unknown whether sigmavirus paralysis occurs in larval stages. Conversely, sigmavirus-induced CO_2_ paralysis and mortality could pose challenges in mass-rearing of fruit flies for SIT, if CO_2_ is used at cool temperatures at any stage of fly processing. Furthermore, our study provides a simple basis for the development of selection procedures for sigmavirus-free individuals.

Similarly, restricted ventilation or CO_2_ treatment at low temperatures has been suggested as treatment option of honey bee colonies infested with *Varroa* during winter, as it was found that this increases *Varroa* mortality (Kozak and Currie 2011; Bahreini and Currie 2015; Onayemi et al. 2022). The presence of sigmavirus or other rhabdoviruses in the studied honey bee colonies and *Varroa* populations, however, is unknown, yet could have contributed to the variability of measured effects.

### Host effects and virus competition

We do not know if BtSV effects on host fitness parameters other than the CO_2_-triggered paralysis and mortality, such as the effects on fertility, fecundity, embryo viability, hatch rate, developmental time, growth rates, lifespan and immunity observed for DmelSV in *D. melanogaster* (Seecof 1964; Fleuriet 1981a) also apply to *B. tryoni*. A previous comprehensive field survey detected BtSV in 22 of 29 field populations (76%) at an overall low prevalence (13.7%) in individuals (Sharpe et al. 2024). In contrast, our study showed that BtSV occurred in 50% of tested laboratory populations and reached high prevalence (80%) in some of these. This indicates that there are environmental conditions under which BtSV is either costly to the host or BtSV does not replicate well. It could also be that host genotypic differences affect virus transmission efficiency as demonstrated for *Drosophila* (Wilfert and Jiggins 2010). Furthermore, our study confirmed that the previously detected negative association pattern between BtSV and BtIV in field populations (Sharpe et al. 2024) may indeed be due to their mostly maternal transmission which would reduce chances of the two vertically transmitted viruses to coinfect the same maternal lineages. The limited efficiency of paternal and horizontal transmission of BtSV may prevent the establishment of BtSV and BtIV coinfections in *B. tryoni*, possibly in addition to competitive exclusion effects such as immune priming or direct competition.

## CONCLUSIONS

Virus discovery from insect transcriptomes revealed virome diversity in tephritid fruit flies, including a diversity of sigmaviruses (Longdon et al. 2017; Zhang et al. 2020; Sharpe et al. 2021; Pradhan et al. 2024a) whereby BtSV is structurally different from DMelSV and other sigmaviruses because it does not have the X ORF. Our results revealed asymmetric patterns of biparental BtSV transmission in *B. tryoni*, with efficient maternal, limited paternal and inefficient horizontal transmission, as well as CO_2_-triggered paralysis and mortality. These findings parallel sigmavirus research in *Drosophila* and other hosts, revealing general features of sigmavirus and rhabdovirus interactions across diverse host species. Future studies on host-virus interactions could enhance existing pest management strategies such as SIT and develop novel pest management strategies using viruses as biocontrol agent or target for CO_2_ disinfestation treatments.

## ACKNOWLEDGEMENTS

We thank Carl Ramirez for help with the establishment of the BtCHome21, BtCS8.17 and BtCEC16. We thank Stephen Sharpe for help with the Rotorgene robotics for RT-qPCR analysis.

## AUTHOR CONTRIBUTIONS

The study was conceived and designed by SKP, MR and JLM, as part of the PhD project of SKP supervised by MR, JLM, SB and AR. Data collection was performed by SKP with the help of GT and FB, and analyses were carried out by SKP and JLM. SKP, MR and JLM wrote the manuscript. All authors contributed to the final draft and approved the manuscript for submission.

## FUNDING

The research was supported by The Fresh and Secure Trade Alliance funded through the Hort Frontiers International Markets Fund, part of the Hort Frontiers strategic partnership initiative developed by Hort Innovation, with co-investment from the Queensland Department of Agriculture and Fisheries (Queensland), Department of Primary Industries and Regional Development (South Australia), Department of Energy, Environment and Climate Action (Victoria), Department of Tourism, Industry and Trade, Department of Primary Industries and Regions (Northern Territory), Department of Natural Resources and Environment (NSW), Queensland University of Technology, James Cook University, Western Sydney University, Australian Blueberry Growers’ Association, GreenSkin Avocados, and contributions from the Australian Government and the strawberry and avocado R&D levy. SKP was supported by an ICAR Senior Research Fellowship and a WSU PhD scholarship award.

## COMPETING INTEREST

The authors declare that they do not have a conflict of interest.

## Notes

### Competing Interest Statement

The authors have declared no competing interest.

