## Supplementary Information for "Asymmetric biparental but inefficient horizontal transmission of paralysis-causing sigmavirus in Queensland fruit fly"

### Supplementary Information (SI)

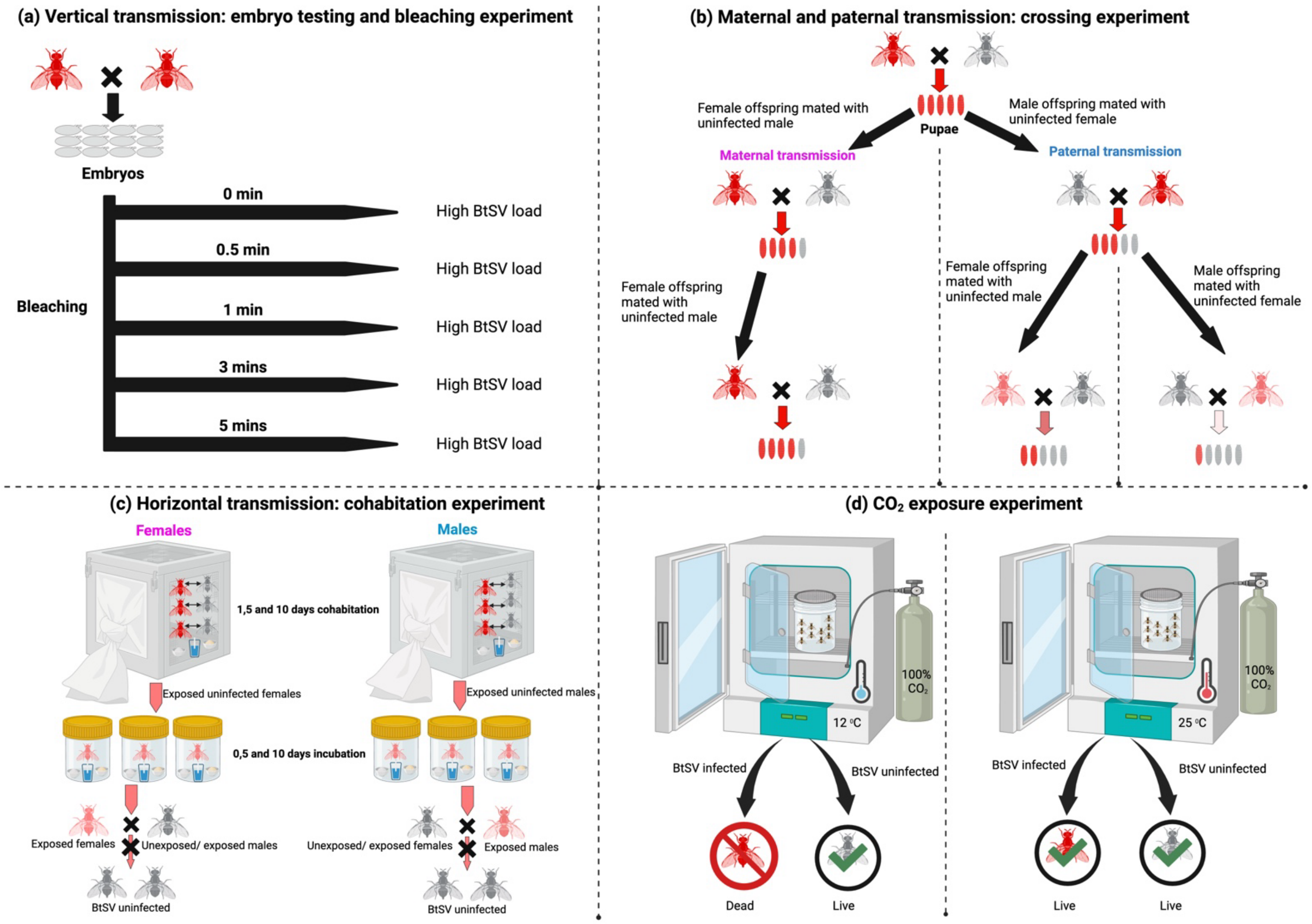


**SI 1. Overview of the experiments performed to assess transmission pathways and host effects of Bactrocera tryoni sigmavirus (BtSV).** Experiments to assess **(a)** vertical transmission of BtSV within embryos (embryo testing and bleaching experiments), **(b)** maternal and paternal BtSV transmission (crossing experiments), **(c)** horizontal BtSV transmission (cohabitation experiments) and **(d)** $CO_2$-triggered paralysis ($CO_2$ exposure experiment).

### SI 2. Nucleotide sequence of sigmavirus variant (BtSV1) found in C28+ and LE+

>C28+.Female1.SV1
GCTATCGGGTGGTTTGTGGAGTCTGACTCGTGTTTGGCTCATGCCATTTATGATAACTTAAAGGCCCTAACAGGTGAGGA
CTGGAGTAATTCAGTTGAAGGATTCAAACGAACTGGGTCAGCATTACACAGATTTTCTTGTTCGAGGCAAAGTGCTGGAG
GGTTTATTGCATCTAGTCCAATCAAGTCGACGTATATGATATCAACAACAAACACATTATCTGAGCTCGGTGATAAAAACT
ATGATTTTATTTTCCAAACATTGTTGCTCTACACTGAAATCACTTGCGGAGAATTACATGCTAATAACCCGAACTCAGGATA
TTACCATATGCATATCTCATGTTCAAATTGTTTACGAGAAATTGAGGAACCAATATTGAATACCTCTGTGACGCCAACCTAT
CGT
>C28+.Male1.SV1
GCTATCGGGTGGTTTGTGGAGTCTGACTCGTGTTTGGCTCATGCCATTTATGATAACTTAAAGGCCCTAACAGGTGAGGA
CTGGAGTAATTCAGTTGAAGGATTCAAACGAACTGGGTCAGCATTACACAGATTTTCTTGTTCGAGGCAAAGTGCTGGAG
GGTTTATTGCATCTAGTCCAATCAAGTCGACGTATATGATATCAACAACAAACACATTATCTGAGCTCGGTGATAAAAACT
ATGATTTTATTTTCCAAACATTGTTGCTCTACACTGAAATCACTTGCGGAGAATTACATGCTAATAACCCGAACTCAGGATA
TTACCATATGCATATCTCATGTTCAAATTGTTTACGAGAAATTGAGGAACCAATATTGAATACCTCTGTGACGCCAACCTAT
CGT
>C28+.Male2.SV1
GCTATCGGGTGGTTTGTGGAGTCTGACTCGTGTTTGGCTCATGCCATTTATGATAACTTAAAGGCCCTAACAGGTGAGGA
CTGGAGTAATTCAGTTGAAGGATTCAAACGAACTGGGTCAGCATTACACAGATTTTCTTGTTCGAGGCAAAGTGCTGGAG
GGTTTATTGCATCTAGTCCAATCAAGTCGACGTATATGATATCAACAACAAACACATTATCTGAGCTCGGTGATAAAAACT
ATGATTTTATTTTCCAAACATTGTTGCTCTACACTGAAATCACTTGCGGAGAATTACATGCTAATAACCCGAACTCAGGATA
TTACCATATGCATATCTCATGTTCAAATTGTTTACGAGAAATTGAGGAACCAATATTGAATACCTCTGTGACGCCAACCTAT
CGT
>LE+.Female1.SV1
GCTATCGGGTGGTTTGTGGAGTCTGACTCGTGTTTGGCTCATGCCATTTATGATAACTTAAAGGCCCTAACAGGTGAGGA
CTGGAGTAATTCAGTTGAAGGATTCAAACGAACTGGGTCAGCATTACACAGATTTTCTTGTTCGAGGCAAAGTGCTGGAG
GGTTTATTGCATCTAGTCCAATCAAGTCGACGTATATGATATCAACAACAAACACATTATCTGAGCTCGGTGATAAAAACT
ATGATTTTATTTTCCAAACATTGTTGCTCTACACTGAAATCACTTGCGGAGAATTACATGCTAATAACCCGAACTCAGGATA
TTACCATATGCATATCTCATGTTCAAATTGTTTACGAGAAATTGAGGAACCAATATTGAATACCTCTGTGACGCCAACCTAT
CGT
>LE+.Male1.SV1
GCTATCGGGTGGTTTGTGGAGTCTGACTCGTGTTTGGCTCATGCCATTTATGATAACTTAAAGGCCCTAACAGGTGAGGA
CTGGAGTAATTCAGTTGAAGGATTCAAACGAACTGGGTCAGCATTACACAGATTTTCTTGTTCGAGGCAAAGTGCTGGAG
GGTTTATTGCATCTAGTCCAATCAAGTCGACGTATATGATATCAACAACAAACACATTATCTGAGCTCGGTGATAAAAACT
ATGATTTTATTTTCCAAACATTGTTGCTCTACACTGAAATCACTTGCGGAGAATTACATGCTAATAACCCGAACTCAGGATA
TTACCATATGCATATCTCATGTTCAAATTGTTTACGAGAAATTGAGGAACCAATATTGAATACCTCTGTGACGCCAACCTAT
CGT
>LE+.Male2.SV1
GCTATCGGGTGGTTTGTGGAGTCTGACTCGTGTTTGGCTCATGCCATTTATGATAACTTAAAGGCCCTAACAGGTGAGGA
CTGGAGTAATTCAGTTGAAGGATTCAAACGAACTGGGTCAGCATTACACAGATTTTCTTGTTCGAGGCAAAGTGCTGGAG
GGTTTATTGCATCTAGTCCAATCAAGTCGACGTATATGATATCAACAACAAACACATTATCTGAGCTCGGTGATAAAAACT
ATGATTTTATTTTCCAAACATTGTTGCTCTACACTGAAATCACTTGCGGAGAATTACATGCTAATAACCCGAACTCAGGATA
TTACCATATGCATATCTCATGTTCAAATTGTTTACGAGAAATTGAGGAACCAATATTGAATACCTCTGTGACGCCAACCTAT
CGT

### SI 3. BtSV loads across developmental stages

Kruskal-Wallis rank sum test

Kruskal-Wallis chi-squared = 17.99, df = 4, p-value = 0.0012

Dunn (1964) Kruskal-Wallis multiple comparison
  p-values adjusted with the Benjamini-Hochberg method.

| | Comparison | Z | P.unadj | P.adj |
|---|---|---|---|---|
| 1 | Embryos - Female | -2.52435736 | 0.0115910033 | 0.038636678 |
| 2 | Embryos - Larvae | -1.70576206 | 0.0880523901 | 0.176104780 |
| 3 | Female - Larvae | 1.13160847 | 0.2577990769 | 0.368284396 |
| 4 | Embryos - Male | -3.06839989 | 0.0021520842 | 0.010760421 |
| 5 | Female - Male | -0.47115466 | 0.6375302971 | 0.708366997 |
| 6 | Larvae - Male | -1.67565101 | 0.0938065718 | 0.156344286 |
| 7 | Embryos - Pupae | -3.71802824 | 0.0002007838 | 0.002007838 |
| 8 | Female - Pupae | -0.51139998 | 0.6090710073 | 0.761338759 |
| 9 | Larvae - Pupae | -2.01226618 | 0.0441918858 | 0.110479714 |
| 10 | Male - Pupae | 0.03264255 | 0.9739596363 | 0.973959636 |

### SI 4. Test for transovarian or transovum transmission of BtSV in LE+ embryos

Kruskal-Wallis rank sum test

Kruskal-Wallis chi-squared = 9.539, df = 4, p-value = 0.04895

Dunn (1964) Kruskal-Wallis multiple comparison
  p-values adjusted with the Benjamini-Hochberg method.

| | Comparison | Z | P.unadj | P.adj |
|---|---|---|---|---|
| 1 | 1min - 3min | 1.10452021 | 0.269367543 | 0.44894591 |
| 2 | 1min - 5min | -0.44871133 | 0.653639908 | 0.72626656 |
| 3 | 3min - 5min | -1.55323154 | 0.120367829 | 0.24073566 |
| 4 | 1min - Female | -0.55677679 | 0.577679940 | 0.72209993 |
| 5 | 3min - Female | -1.88887828 | 0.058908140 | 0.19636047 |
| 6 | 5min - Female | -0.01561056 | 0.987545078 | 0.98754508 |
| 7 | 1min - 0min | 1.72581282 | 0.084381092 | 0.21095273 |
| 8 | 3min - 0min | 0.62129262 | 0.534407111 | 0.76343873 |
| 9 | 5min - 0min | 2.17452416 | 0.029665793 | 0.14832897 |
| 10 | Female - 0min | 2.63818537 | 0.008335101 | 0.08335101 |

**SI 5. Test for transovarian or transovum transmission of BtSV in embryos of (LE$^+$ × B6$^-$) × B6$^-$ and B6$^-$ × (LE$^+$ × B6$^-$) in two families**

### Embryos of (LE$^+$ × B6$^-$) × B6$^-$ family 5

Kruskal-Wallis rank sum test

Kruskal-Wallis chi-squared = 5.4218, df = 5, p-value = 0.3666

Dunn (1964) Kruskal-Wallis multiple comparison
p-values adjusted with the Benjamini-Hochberg method.

| | Comparison | Z | P.unadj | P.adj |
|---|---|---|---|---|
| 1 | 1min - 0.5min | -0.9341987 | 0.35020139 | 0.6566276 |
| 2 | 1min - 3min | -0.4670994 | 0.64042879 | 0.7389563 |
| 3 | 0.5min - 3min | 0.4670994 | 0.64042879 | 0.8005360 |
| 4 | 1min - 5min | 0.4670994 | 0.64042879 | 0.8733120 |
| 5 | 0.5min - 5min | 1.4012981 | 0.16112495 | 0.6042186 |
| 6 | 3min - 5min | 0.9341987 | 0.35020139 | 0.7504315 |
| 7 | 1min - Female | 0.2412091 | 0.80939308 | 0.8093931 |
| 8 | 0.5min - Female | 1.4472545 | 0.14782567 | 0.7391284 |
| 9 | 3min - Female | 0.8442318 | 0.39853991 | 0.6642332 |
| 10 | 5min - Female | -0.3618136 | 0.71749132 | 0.7687407 |
| 11 | 1min - 0min | 1.1677484 | 0.24290826 | 0.6072707 |
| 12 | 0.5min - 0min | 2.1019471 | 0.03555791 | 0.5333686 |
| 13 | 3min - 0min | 1.6348478 | 0.10208096 | 0.7656072 |
| 14 | 5min - 0min | 0.7006490 | 0.48352206 | 0.7252831 |
| 15 | Female - 0min | 1.2663476 | 0.20538865 | 0.6161659 |

Kruskal-Wallis rank sum test when compared all embryos into one pool with females
Kruskal-Wallis chi-squared = 0.098182, df = 1, p-value = 0.754

### Embryos of (LE$^+$ × B6$^-$) × B6$^-$ family 6

Kruskal-Wallis rank sum test

Kruskal-Wallis chi-squared = 2.1491, df = 5, p-value = 0.8282

Dunn (1964) Kruskal-Wallis multiple comparison
p-values adjusted with the Benjamini-Hochberg method.

| | Comparison | Z | P.unadj | P.adj |
|---|---|---|---|---|
| 1 | 1min - 0.5min | 0.46709937 | 0.6404288 | 1.0000000 |
| 2 | 1min - 3min | -0.46709937 | 0.6404288 | 1.0000000 |
| 3 | 0.5min - 3min | -0.93419873 | 0.3502014 | 1.0000000 |
| 4 | 1min - 5min | 0.23354968 | 0.8153346 | 0.8735728 |
| 5 | 0.5min - 5min | -0.23354968 | 0.8153346 | 0.9407707 |
| 6 | 3min - 5min | 0.70064905 | 0.4835221 | 1.0000000 |
| 7 | 1min - Female | 0.66332496 | 0.5071225 | 1.0000000 |
| 8 | 0.5min - Female | 0.06030227 | 0.9519149 | 0.9519149 |
| 9 | 3min - Female | 1.26634765 | 0.2053886 | 1.0000000 |
| 10 | 5min - Female | 0.36181361 | 0.7174913 | 1.0000000 |
| 11 | 1min - 0min | 0.70064905 | 0.4835221 | 1.0000000 |
| 12 | 0.5min - 0min | 0.23354968 | 0.8153346 | 1.0000000 |
| 13 | 3min - 0min | 1.16774842 | 0.2429083 | 1.0000000 |
| 14 | 5min - 0min | 0.46709937 | 0.6404288 | 1.0000000 |
| 15 | Female - 0min | 0.24120908 | 0.8093931 | 1.0000000 |

Kruskal-Wallis rank sum test when compared all embryos into one pool with females
Kruskal-Wallis chi-squared = 0.53455, df = 1, p-value = 0.4647

**Embryos of B6$^-$ × (LE$^+$ × B6$^-$) family 5**

Kruskal-Wallis rank sum test

Kruskal-Wallis chi-squared = 7.3333, df = 5, p-value = 0.197

Dunn (1964) Kruskal-Wallis multiple comparison
p-values adjusted with the Benjamini-Hochberg method.

| | Comparison | Z | P.unadj | P.adj |
|---|---|---|---|---|
| 1 | 1min - 0.5min | -0.2581989 | 0.79625341 | 0.7962534 |
| 2 | 1min - 3min | -0.5163978 | 0.60557662 | 0.8257863 |
| 3 | 0.5min - 3min | -0.2581989 | 0.79625341 | 0.8531287 |
| 4 | 1min - 5min | 0.2581989 | 0.79625341 | 0.9187539 |
| 5 | 0.5min - 5min | 0.5163978 | 0.60557662 | 0.9083649 |
| 6 | 3min - 5min | 0.7745967 | 0.43857803 | 0.8223338 |
| 7 | 1min - Male | -1.4696938 | 0.14164469 | 0.7082235 |
| 8 | 0.5min - Male | -1.1430952 | 0.25299906 | 0.9487465 |
| 9 | 3min - Male | -0.8164966 | 0.41421618 | 1.0000000 |
| 10 | 5min - Male | -1.7962925 | 0.07244801 | 0.5433601 |
| 11 | 1min - 0min | 0.5163978 | 0.60557662 | 1.0000000 |
| 12 | 0.5min - 0min | 0.7745967 | 0.43857803 | 0.9398101 |
| 13 | 3min - 0min | 1.0327956 | 0.30169958 | 0.9050987 |
| 14 | 5min - 0min | 0.2581989 | 0.79625341 | 0.9953168 |
| 15 | Male - 0min | 2.1228911 | 0.03376298 | 0.5064447 |

Kruskal-Wallis rank sum test when compared all embryos into one pool with males

Kruskal-Wallis chi-squared = 6, df = 1, p-value = 0.01431

**Embryos of B6$^-$ × (LE$^+$ × B6$^-$) family 6**

Kruskal-Wallis rank sum test
Kruskal-Wallis chi-squared = 7.9091, df = 5, p-value = 0.1613

Dunn (1964) Kruskal-Wallis multiple comparison
p-values adjusted with the Benjamini-Hochberg method.

| | Comparison | Z | P.unadj | P.adj |
|---|---|---|---|---|
| 1 | 1min - 0.5min | 0.2335497 | 0.81533459 | 0.8153346 |
| 2 | 1min - 3min | -0.4670994 | 0.64042879 | 0.8733120 |
| 3 | 0.5min - 3min | -0.7006490 | 0.48352206 | 0.9066039 |
| 4 | 1min - 5min | -0.2335497 | 0.81533459 | 0.8735728 |
| 5 | 0.5min - 5min | -0.4670994 | 0.64042879 | 0.9606432 |
| 6 | 3min - 5min | 0.2335497 | 0.81533459 | 0.9407707 |
| 7 | 1min - Male | -1.5075567 | 0.13166802 | 0.6583401 |
| 8 | 0.5min - Male | -1.8090681 | 0.07044043 | 0.5283032 |
| 9 | 3min - Male | -0.9045340 | 0.36571230 | 0.9142807 |
| 10 | 5min - Male | -1.2060454 | 0.22779999 | 0.8542500 |
| 11 | 1min - 0min | 0.4670994 | 0.64042879 | 1.0000000 |
| 12 | 0.5min - 0min | 0.2335497 | 0.81533459 | 1.0000000 |
| 13 | 3min - 0min | 0.9341987 | 0.35020139 | 1.0000000 |
| 14 | 5min - 0min | 0.7006490 | 0.48352206 | 1.0000000 |
| 15 | Male - 0min | 2.1105794 | 0.03480848 | 0.5221272 |

Kruskal-Wallis rank sum test when compared all embryos into one pool with males
Kruskal-Wallis chi-squared = 6.8182, df = 1, p-value = 0.009023

### SI. 6. Horizontal transmission of BtSV

### Horizontal transmission of BtSV in female

```
Kruskal-Wallis rank sum test

Kruskal-Wallis chi-squared = 31.263, df = 10, p-value = 0.0005308

Dunn (1964) Kruskal-Wallis multiple comparison
  p-values adjusted with the Benjamini-Hochberg method.
```

| | Comparison | Z | P.unadj | P.adj |
|---|---|---|---|---|
| 1 | 1:0 - 1:10 | 2.629213943 | 8.558250e-03 | 0.0588379689 |
| 2 | 1:0 - 1:5 | 0.553518725 | 5.799083e-01 | 0.7248853279 |
| 3 | 1:10 - 1:5 | -2.075695218 | 3.792215e-02 | 0.1390478778 |
| 4 | 1:0 - 10:0 | 0.805593257 | 4.204774e-01 | 0.6423960501 |
| 5 | 1:10 - 10:0 | -1.471372809 | 1.411903e-01 | 0.2986718365 |
| 6 | 1:5 - 10:0 | 0.326231980 | 7.442488e-01 | 0.8709295018 |
| 7 | 1:0 - 10:10 | 0.572570414 | 5.669356e-01 | 0.7251501816 |
| 8 | 1:10 - 10:10 | -1.704395652 | 8.830720e-02 | 0.2312807505 |
| 9 | 1:5 - 10:10 | 0.093209137 | 9.257374e-01 | 0.9791453434 |
| 10 | 10:0 - 10:10 | -0.208421967 | 8.348995e-01 | 0.9003818199 |
| 11 | 1:0 - 10:5 | -0.226365048 | 8.209175e-01 | 0.9030092615 |
| 12 | 1:10 - 10:5 | -2.503331114 | 1.230304e-02 | 0.0751852336 |
| 13 | 1:5 - 10:5 | -0.705726325 | 4.803583e-01 | 0.6443831482 |
| 14 | 10:0 - 10:5 | -0.923011568 | 3.560012e-01 | 0.6118770227 |
| 15 | 10:10 - 10:5 | -0.714589601 | 4.748627e-01 | 0.6529361744 |
| 16 | 1:0 - 5:0 | -0.253696082 | 7.997304e-01 | 0.9163577053 |
| 17 | 1:10 - 5:0 | -2.882910025 | 3.940200e-03 | 0.0361184971 |
| 18 | 1:5 - 5:0 | -0.807214807 | 4.195427e-01 | 0.6592814428 |
| 19 | 10:0 - 5:0 | -1.025300509 | 3.052214e-01 | 0.5595725950 |
| 20 | 10:10 - 5:0 | -0.792277666 | 4.281988e-01 | 0.6365117165 |
| 21 | 10:5 - 5:0 | 0.006657796 | 9.946879e-01 | 0.9946878870 |
| 22 | 1:0 - 5:10 | 0.645771846 | 5.184271e-01 | 0.6788926623 |
| 23 | 1:10 - 5:10 | -1.983442097 | 4.731807e-02 | 0.1445829879 |
| 24 | 1:5 - 5:10 | 0.092253121 | 9.264969e-01 | 0.9614590821 |
| 25 | 10:0 - 5:10 | -0.246338434 | 8.054203e-01 | 0.9040431493 |
| 26 | 10:10 - 5:10 | -0.013315591 | 9.893760e-01 | 1.0000000000 |
| 27 | 10:5 - 5:10 | 0.785619871 | 4.320902e-01 | 0.6253937388 |
| 28 | 5:0 - 5:10 | 0.899467928 | 3.684035e-01 | 0.6140057860 |
| 29 | 1:0 - 5:5 | 1.499113213 | 1.338443e-01 | 0.3067264392 |
| 30 | 1:10 - 5:5 | -1.130100730 | 2.584338e-01 | 0.4901330353 |
| 31 | 1:5 - 5:5 | 0.945594488 | 3.443555e-01 | 0.6109532294 |
| 32 | 10:0 - 5:5 | 0.492676868 | 6.222409e-01 | 0.7605166857 |
| 33 | 10:10 - 5:5 | 0.725699711 | 4.680229e-01 | 0.6600322650 |
| 34 | 10:5 - 5:5 | 1.524635173 | 1.273501e-01 | 0.3045328939 |
| 35 | 5:0 - 5:5 | 1.752809295 | 7.963475e-02 | 0.2189955545 |
| 36 | 5:10 - 5:5 | 0.853341367 | 3.934700e-01 | 0.6364956303 |
| 37 | 1:0 - Infected | -1.641692317 | 1.006538e-01 | 0.2516344616 |
| 38 | 1:10 - Infected | -4.120538995 | 3.779871e-05 | 0.0010394645 |
| 39 | 1:5 - Infected | -2.163554776 | 3.049854e-02 | 0.1198156754 |
| 40 | 10:0 - Infected | -2.212209374 | 2.695220e-02 | 0.1140285446 |
| 41 | 10:10 - Infected | -1.989396919 | 4.665741e-02 | 0.1509504424 |
| 42 | 10:5 - Infected | -1.225468502 | 2.203988e-01 | 0.4329261175 |
| 43 | 5:0 - Infected | -1.402505357 | 1.607644e-01 | 0.3274830206 |
| 44 | 5:10 - Infected | -2.250531852 | 2.441520e-02 | 0.1119030179 |
| 45 | 5:5 - Infected | -3.055069809 | 2.250081e-03 | 0.0309386196 |
| 46 | 1:0 - Uninfected | 2.897423858 | 3.762411e-03 | 0.0413865161 |
| 47 | 1:10 - Uninfected | 0.418577180 | 6.755252e-01 | 0.8076931379 |
| 48 | 1:5 - Uninfected | 2.375561399 | 1.752228e-02 | 0.0963725231 |
| 49 | 10:0 - Uninfected | 1.774542052 | 7.597353e-02 | 0.2199233843 |
| 50 | 10:10 - Uninfected | 1.997354507 | 4.578669e-02 | 0.1573917335 |
| 51 | 10:5 - Uninfected | 2.761282924 | 5.757478e-03 | 0.0452373244 |
| 52 | 5:0 - Uninfected | 3.136610818 | 1.709128e-03 | 0.0313340193 |
| 53 | 5:10 - Uninfected | 2.288584323 | 2.210352e-02 | 0.1105175781 |
| 54 | 5:5 - Uninfected | 1.484046366 | 1.377966e-01 | 0.3031525648 |
| 55 | Infected - Uninfected | 4.306183703 | 1.660951e-05 | 0.0009135233 |

**Horizontal transmission of BtSV in male**

Kruskal-Wallis rank sum test
Kruskal-Wallis chi-squared = 28.936, df = 10, p-value = 0.001276
Dunn (1964) Kruskal-Wallis multiple comparison
p-values adjusted with the Benjamini-Hochberg method.

| | Comparison | Z | P.unadj | P.adj |
|---|---|---|---|---|
| 1 | 1:0 - 1:10 | 1.7064938 | 8.791617e-02 | 0.268632751 |
| 2 | 1:0 - 1:5 | -0.2201928 | 8.257210e-01 | 0.908293144 |
| 3 | 1:10 - 1:5 | -1.9266866 | 5.401870e-02 | 0.198068567 |
| 4 | 1:0 - 10:0 | -1.1284879 | 2.591139e-01 | 0.431856569 |
| 5 | 1:10 - 10:0 | -2.8349817 | 4.582831e-03 | 0.063013928 |
| 6 | 1:5 - 10:0 | -0.9082951 | 3.637223e-01 | 0.540668324 |
| 7 | 1:0 - 10:10 | -0.8257228 | 4.089613e-01 | 0.591917732 |
| 8 | 1:10 - 10:10 | -2.5322167 | 1.133439e-02 | 0.069265736 |
| 9 | 1:5 - 10:10 | -0.6055301 | 5.448269e-01 | 0.713463733 |
| 10 | 10:0 - 10:10 | 0.3027650 | 7.620689e-01 | 0.855383494 |
| 11 | 1:0 - 10:5 | -0.6605783 | 5.088828e-01 | 0.682647679 |
| 12 | 1:10 - 10:5 | -2.3670721 | 1.792944e-02 | 0.082176599 |
| 13 | 1:5 - 10:5 | -0.4403855 | 6.596579e-01 | 0.788721427 |
| 14 | 10:0 - 10:5 | 0.4679096 | 6.398492e-01 | 0.782037957 |
| 15 | 10:10 - 10:5 | 0.1651446 | 8.688302e-01 | 0.918955016 |
| 16 | 1:0 - 5:0 | -0.9908674 | 3.217503e-01 | 0.491563017 |
| 17 | 1:10 - 5:0 | -2.6973612 | 6.989141e-03 | 0.054914679 |
| 18 | 1:5 - 5:0 | -0.7706746 | 4.408998e-01 | 0.621781783 |
| 19 | 10:0 - 5:0 | 0.1376205 | 8.905404e-01 | 0.907031865 |
| 20 | 10:10 - 5:0 | -0.1651446 | 8.688302e-01 | 0.936973742 |
| 21 | 10:5 - 5:0 | -0.3302891 | 7.411815e-01 | 0.849270475 |
| 22 | 1:0 - 5:10 | 0.3578132 | 7.204831e-01 | 0.843118514 |
| 23 | 1:10 - 5:10 | -1.3486806 | 1.774396e-01 | 0.375352948 |
| 24 | 1:5 - 5:10 | 0.5780060 | 5.632601e-01 | 0.720448945 |
| 25 | 10:0 - 5:10 | 1.4863011 | 1.371995e-01 | 0.342998751 |
| 26 | 10:10 - 5:10 | 1.1835360 | 2.365968e-01 | 0.419768461 |
| 27 | 10:5 - 5:10 | 1.0183915 | 3.084919e-01 | 0.484773062 |
| 28 | 5:0 - 5:10 | 1.3486806 | 1.774396e-01 | 0.390367066 |
| 29 | 1:0 - 5:5 | 0.4679096 | 6.398492e-01 | 0.799811547 |
| 30 | 1:10 - 5:5 | -1.2385842 | 2.154995e-01 | 0.408705965 |
| 31 | 1:5 - 5:5 | 0.6881024 | 4.913883e-01 | 0.675658952 |
| 32 | 10:0 - 5:5 | 1.5963975 | 1.104001e-01 | 0.319579188 |
| 33 | 10:10 - 5:5 | 1.2936324 | 1.957924e-01 | 0.398836405 |
| 34 | 10:5 - 5:5 | 1.1284879 | 2.591139e-01 | 0.445352087 |
| 35 | 5:0 - 5:5 | 1.4587770 | 1.446265e-01 | 0.345845974 |
| 36 | 5:10 - 5:5 | 0.1100964 | 9.123329e-01 | 0.912332942 |
| 37 | 1:0 - Infected | -2.3945962 | 1.663868e-02 | 0.083193393 |
| 38 | 1:10 - Infected | -4.1010900 | 4.112085e-05 | 0.002261647 |
| 39 | 1:5 - Infected | -2.1744034 | 2.967485e-02 | 0.116579769 |
| 40 | 10:0 - Infected | -1.2661083 | 2.054743e-01 | 0.403610234 |
| 41 | 10:10 - Infected | -1.5688734 | 1.166774e-01 | 0.305583795 |
| 42 | 10:5 - Infected | -1.7340179 | 8.291490e-02 | 0.268254094 |
| 43 | 5:0 - Infected | -1.4037288 | 1.603996e-01 | 0.367582469 |
| 44 | 5:10 - Infected | -2.7524094 | 5.915851e-03 | 0.065074362 |
| 45 | 5:5 - Infected | -2.8625058 | 4.203056e-03 | 0.077056023 |
| 46 | 1:0 - Uninfected | 1.5688734 | 1.166774e-01 | 0.320862985 |
| 47 | 1:10 - Uninfected | -0.1376205 | 8.905404e-01 | 0.924145673 |
| 48 | 1:5 - Uninfected | 1.7890661 | 7.360417e-02 | 0.253014327 |
| 49 | 10:0 - Uninfected | 2.6973612 | 6.989141e-03 | 0.064067125 |
| 50 | 10:10 - Uninfected | 2.3945962 | 1.663868e-02 | 0.091512732 |
| 51 | 10:5 - Uninfected | 2.2294516 | 2.578387e-02 | 0.109085606 |
| 52 | 5:0 - Uninfected | 2.5597408 | 1.047503e-02 | 0.072015812 |
| 53 | 5:10 - Uninfected | 1.2110601 | 2.258724e-01 | 0.414099319 |
| 54 | 5:5 - Uninfected | 1.1009638 | 2.709124e-01 | 0.438240692 |
| 55 | Infected - Uninfected | 3.9634696 | 7.386829e-05 | 0.002031378 |

**Comparison of acquired BtSV load between females and males of EC16$^{-}$**

Kruskal-Wallis rank sum test
Kruskal-Wallis chi-squared = 1.141, df = 1, p-value = 0.2854

**SI 7. Body region and tissue localisation of BtSV in LE+ females and males and in female and male offspring of LE+ × B6–.**

**BtSV tissue localisation in LE+ female body region and tissues**

Kruskal-Wallis rank sum test

Kruskal-Wallis chi-squared = 1.395, df = 2, p-value = 0.4978

Dunn (1964) Kruskal-Wallis multiple comparison
p-values adjusted with the Benjamini-Hochberg method.

| | Comparison | Z | P.unadj | P.adj |
|---|---|---|---|---|
| 1 | Gut - Head | -1.1667262 | 0.2433210 | 0.7299629 |
| 2 | Gut - Ovaries | -0.4242641 | 0.6713732 | 0.6713732 |
| 3 | Head - Ovaries | 0.7424621 | 0.4578074 | 0.6867111 |

**BtSV tissue localisation in LE+ male body region and tissues**

Kruskal-Wallis rank sum test

Kruskal-Wallis chi-squared = 0.26, df = 2, p-value = 0.8781

Dunn (1964) Kruskal-Wallis multiple comparison
p-values adjusted with the Benjamini-Hochberg method.

| | Comparison | Z | P.unadj | P.adj |
|---|---|---|---|---|
| 1 | Gut - Head | -0.1414214 | 0.8875371 | 0.8875371 |
| 2 | Gut - Testes | -0.4949747 | 0.6206179 | 1.0000000 |
| 3 | Head - Testes | -0.3535534 | 0.7236736 | 1.0000000 |

**BtSV tissue localisation in female offspring of LE+ × B6– family 5**

Kruskal-Wallis rank sum test

Kruskal-Wallis chi-squared = 1.8846, df = 2, p-value = 0.3897

Dunn (1964) Kruskal-Wallis multiple comparison
p-values adjusted with the Benjamini-Hochberg method.

| | Comparison | Z | P.unadj | P.adj |
|---|---|---|---|---|
| 1 | Gut - Head | -0.1961161 | 0.8445193 | 0.8445193 |
| 2 | Gut - Ovaries | -1.2747549 | 0.2023960 | 0.6071880 |
| 3 | Head - Ovaries | -1.0786387 | 0.2807488 | 0.4211232 |

**BtSV tissue localisation in female offspring of LE+ × B6– family 6**

Kruskal-Wallis rank sum test

Kruskal-Wallis chi-squared = 9.8462, df = 2, p-value = 0.007277

Dunn (1964) Kruskal-Wallis multiple comparison
p-values adjusted with the Benjamini-Hochberg method.

| | Comparison | Z | P.unadj | P.adj |
|---|---|---|---|---|
| 1 | Gut - Head | -1.568929 | 0.116664465 | 0.116664465 |
| 2 | Gut - Ovaries | -3.137858 | 0.001701872 | 0.005105616 |
| 3 | Head - Ovaries | -1.568929 | 0.116664465 | 0.174996697 |

### BtSV tissue localisation in male offspring of LE+ × B6– family 5

```
Kruskal-Wallis rank sum test

Kruskal-Wallis chi-squared = 6.2692, df = 2, p-value = 0.04352

Dunn (1964) Kruskal-Wallis multiple comparison
  p-values adjusted with the Benjamini-Hochberg method.
```

| | Comparison | Z | P.unadj | P.adj |
|---|---|---|---|---|
| 1 | Gut - Head | -1.6669871 | 0.09551696 | 0.14327544 |
| 2 | Gut - Testes | 0.7844645 | 0.43276758 | 0.43276758 |
| 3 | Head - Testes | 2.4514517 | 0.01422813 | 0.04268439 |

### BtSV tissue localisation in male offspring of LE+ × B6– family 6

```
Kruskal-Wallis rank sum test

Kruskal-Wallis chi-squared = 7.7308, df = 2, p-value = 0.02095

Dunn (1964) Kruskal-Wallis multiple comparison
  p-values adjusted with the Benjamini-Hochberg method.
```

| | Comparison | Z | P.unadj | P.adj |
|---|---|---|---|---|
| 1 | Gut - Head | -2.6475678 | 0.00810731 | 0.02432193 |
| 2 | Gut - Testes | -0.5883484 | 0.55629846 | 0.55629846 |
| 3 | Head - Testes | 2.0592194 | 0.03947322 | 0.05920984 |

### BtSV tissue localisation in female offspring of (LE+ × B6–) × B6– family 1

```
Kruskal-Wallis rank sum test

Kruskal-Wallis chi-squared = 4.6222, df = 2, p-value = 0.09915

Dunn (1964) Kruskal-Wallis multiple comparison
  p-values adjusted with the Benjamini-Hochberg method.
```

| | Comparison | Z | P.unadj | P.adj |
|---|---|---|---|---|
| 1 | Gut - Head | 1.4907120 | 0.13603713 | 0.2040557 |
| 2 | Gut - Ovaries | 2.0869968 | 0.03688843 | 0.1106653 |
| 3 | Head - Ovaries | 0.5962848 | 0.55098499 | 0.5509850 |

### BtSV tissue localisation in female offspring of (LE+ × B6–) × B6– family 2

```
Kruskal-Wallis rank sum test

Kruskal-Wallis chi-squared = 4.5714, df = 2, p-value = 0.1017

Dunn (1964) Kruskal-Wallis multiple comparison
  p-values adjusted with the Benjamini-Hochberg method.
```

| | Comparison | Z | P.unadj | P.adj |
|---|---|---|---|---|
| 1 | Gut - Head | 1.069045 | 0.28504941 | 0.28504941 |
| 2 | Gut - Ovaries | 2.138090 | 0.03250944 | 0.09752833 |
| 3 | Head - Ovaries | 1.069045 | 0.28504941 | 0.42757411 |

**BtSV tissue localisation in male offspring of B6– × (LE+ × B6–) family 1**

Kruskal-Wallis rank sum test

Kruskal-Wallis chi-squared = 5.6, df = 2, p-value = 0.06081

Dunn (1964) Kruskal-Wallis multiple comparison
p-values adjusted with the Benjamini-Hochberg method.

| | Comparison | Z | P.unadj | P.adj |
|---|---|---|---|---|
| 1 | Gut - Head | 0.4472136 | 0.65472085 | 0.65472085 |
| 2 | Gut - Testes | 2.2360680 | 0.02534732 | 0.07604196 |
| 3 | Head - Testes | 1.7888544 | 0.07363827 | 0.11045741 |

**BtSV tissue localisation in male offspring of B6– × (LE+ × B6–) family 2**

Kruskal-Wallis rank sum test

Kruskal-Wallis chi-squared = 3.5, df = 2, p-value = 0.1738

Dunn (1964) Kruskal-Wallis multiple comparison
p-values adjusted with the Benjamini-Hochberg method.

| | Comparison | Z | P.unadj | P.adj |
|---|---|---|---|---|
| 1 | Gut - Head | -0.09805807 | 0.92188618 | 0.9218862 |
| 2 | Gut - Testes | 1.56892908 | 0.11666446 | 0.1749967 |
| 3 | Head - Testes | 1.66698715 | 0.09551696 | 0.2865509 |

**BtSV tissue localisation in female offspring of B6– × (LE+ × B6–) family 1**

Kruskal-Wallis rank sum test

Kruskal-Wallis chi-squared = 6.5, df = 2, p-value = 0.03877

Dunn (1964) Kruskal-Wallis multiple comparison
p-values adjusted with the Benjamini-Hochberg method.

| | Comparison | Z | P.unadj | P.adj |
|---|---|---|---|---|
| 1 | Gut - Head | 0.09805807 | 0.92188618 | 0.92188618 |
| 2 | Gut - Ovaries | 2.25533555 | 0.02411227 | 0.07233682 |
| 3 | Head - Ovaries | 2.15727749 | 0.03098405 | 0.04647608 |

**BtSV tissue localisation in female offspring of B6– × (LE+ × B6–) family 2**

Kruskal-Wallis rank sum test

Kruskal-Wallis chi-squared = 7.3846, df = 2, p-value = 0.02491

Dunn (1964) Kruskal-Wallis multiple comparison
p-values adjusted with the Benjamini-Hochberg method.

| | Comparison | Z | P.unadj | P.adj |
|---|---|---|---|---|
| 1 | Gut - Head | 0.000000 | 1.00000000 | 1.00000000 |
| 2 | Gut - Ovaries | 2.353394 | 0.01860293 | 0.02790439 |
| 3 | Head - Ovaries | 2.353394 | 0.01860293 | 0.05580879 |

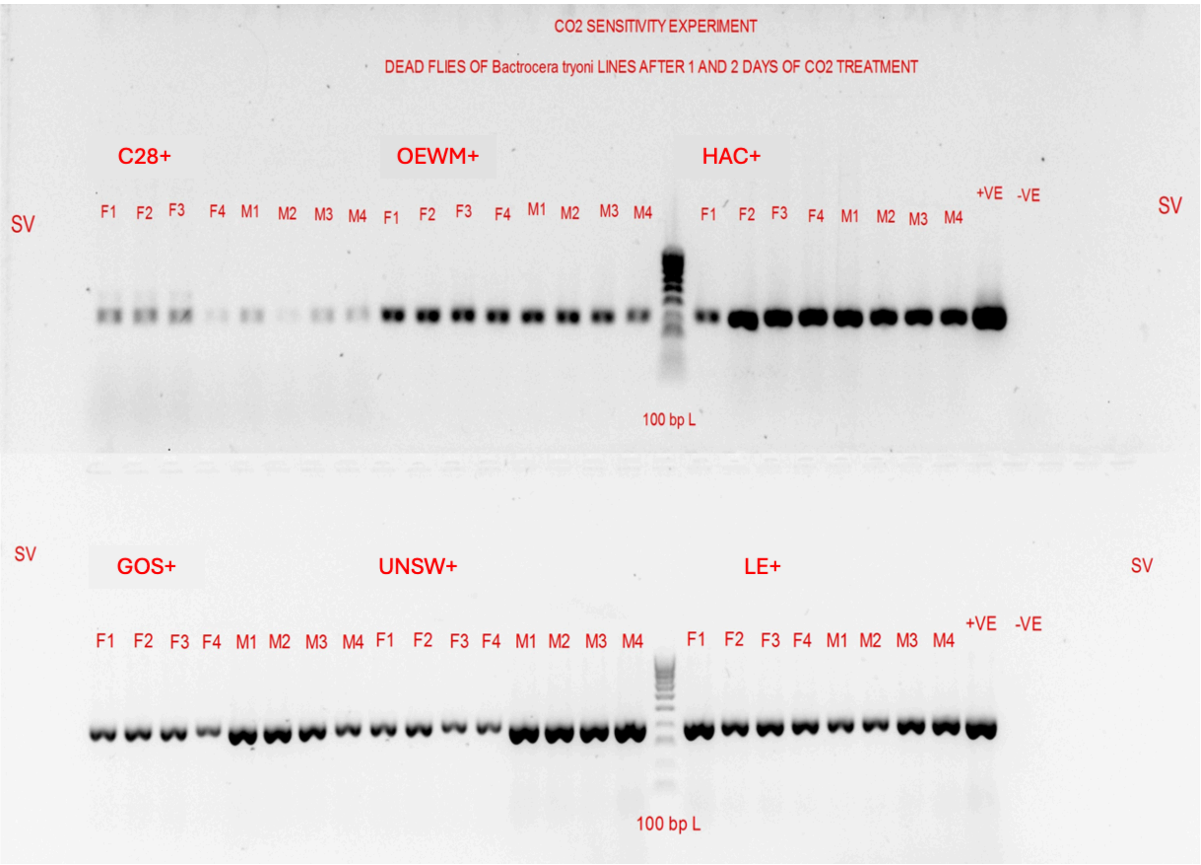


**SI 8.** Standard RT-PCR for the RdRp gene of BtSV in flies that died 1 and 2 days after $CO_2$ treatment of 12 *Bactrocera tryoni* lines.

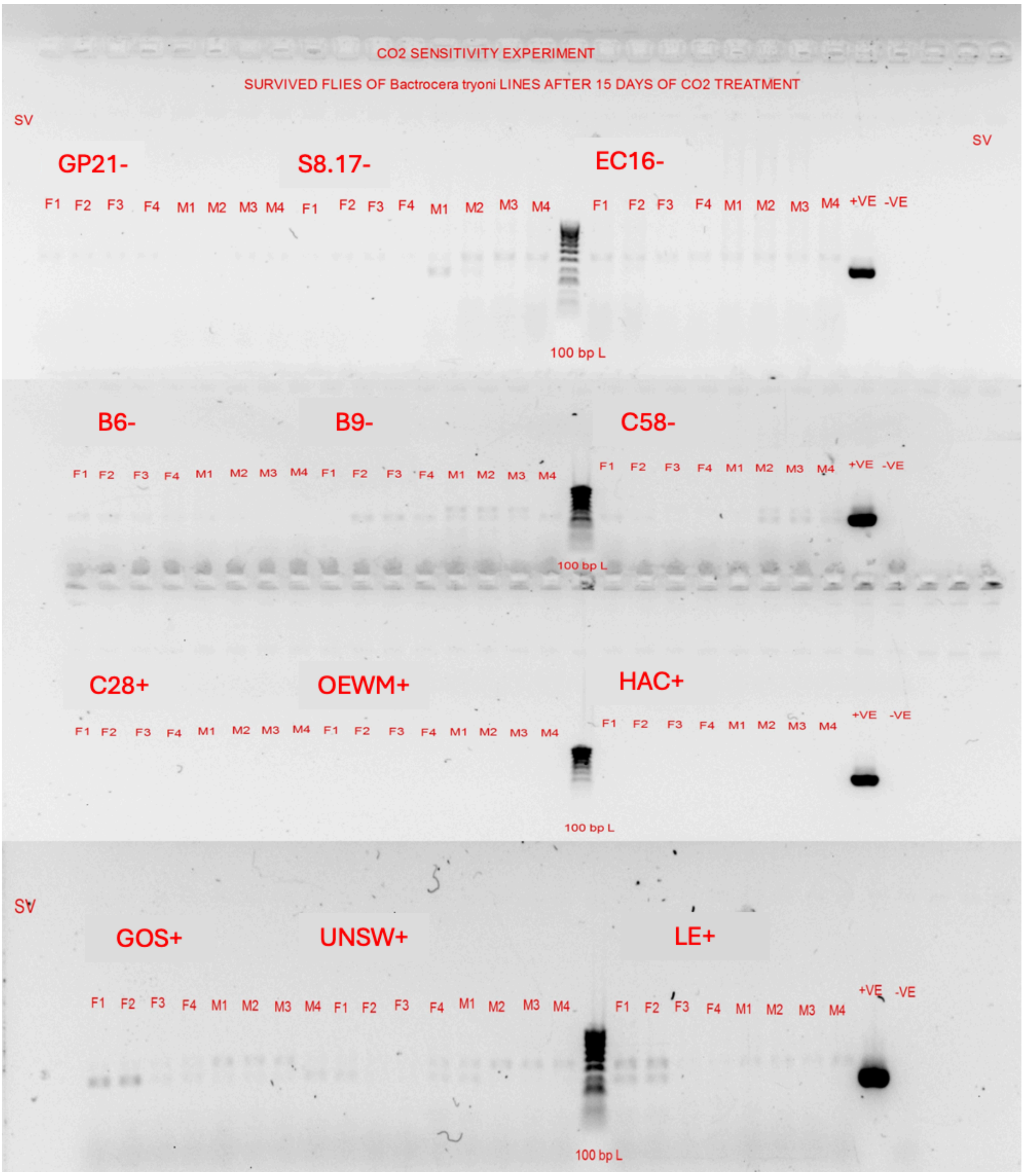


**SI 9.** Standard RT-PCR for the RdRp gene of BtSV in flies that were recovered alive 15 days after $CO_2$ treatment of 12 *Bactrocera tryoni* lines.

**SI 10. Cox proportional hazards model survival analysis of BtSV-infected and uninfected populations**

```
coxph(formula = Surv(Day, Dead > 0) ~ Fly_virus_status, data = data,
    cluster = Population)

  n= 192, number of events= 120
```

| | coef | exp(coef) | se(coef) | robust se | z |
|---|---|---|---|---|---|
| Sigmavirus positive | 0.6363 | 1.8894 | 0.1914 | 0.1794 | 3.547 |

```
Pr(>|z|)
```

**Sigmavirus positive  0.00039 ***

```
Signif. codes:  0 '***' 0.001 '**' 0.01 '*' 0.05 '.' 0.1 ' ' 1
```

| | exp(coef) | exp(-coef) | lower.95 | upper.95 |
|---|---|---|---|---|
| Sigmavirus positive | 1.889 | 0.5293 | 1.329 | 2.686 |

| | |
|---|---|
| Concordance= 0.638 | (se = 0.044 ) |
| Likelihood ratio test | = 11.6 on 1 df, p=7e-04 |
| Wald test | = 12.58 on 1 df, p=4e-04 |
| Score (logrank) test | = 11.43 on 1 df, p=7e-04,Robust = 7.5 p=0.006 |

```
  (Note: the likelihood ratio and score tests assume independence of
     observations within a cluster, the Wald and robust score tests do
not).
```

| confidence interval | 2.5 % | 97.5 % |
|---|---|---|
| Sigmavirus positive | 1.329307 | 2.685597 |

> pval 0.0003901423
